# Translational control of fungal gene expression during the wheat-*Fusarium graminearum* interaction

**DOI:** 10.1101/2021.05.25.445713

**Authors:** Udaykumar Kage, Donald M Gardiner, Jiri Stiller, Kemal Kazan

**Affiliations:** Commonwealth Scientific and Industrial Research Organisation (CSIRO), Agriculture and Food, St Lucia, QLD, Australia 4067

**Keywords:** *Fusarium graminearum*, Fusarium head blight, wheat, ribosome profiling, translation, transcription, ribo-seq, RNA-seq

## Abstract

- To date, translational regulation of key genes controlling infection-related processes in fungal pathogens during their interactions with plants has not been studied. Here, we employed ribosome profiling (ribo-seq) to study translational responses and how such responses are coordinated with transcriptional changes in the fungal pathogen *Fusarium graminearum* (*Fg*), which causes Fusarium head blight (FHB), a destructive disease of cereal crops worldwide.
- Transcription and translation were not always coordinated with approximately 22% of *Fg* genes showing a discordant relationship during wheat infection. Nitrite reductase, which we show here as an important component of fungal virulence, is only regulated at the translational level in *Fg*. In addition, more than 1000 new open reading frames (ORFs), many of which are short and highly conserved, were identified in the *Fg* genome.
- Like in higher eukaryotes, translation is controlled by upstream ORFs (uORFs) in *Fg* during infection. Similarly, miRNAs control both transcription and translation in *Fg* during wheat infection. However, Fgdicer2-dependent miRNAs do not have a significant effect on transcriptional gene expression at the global outset.
- The ribo-seq study undertaken here for the first time in any fungal pathogen discovered novel insights about the biology of an important plant pathogen.

## Introduction

*Fusarium graminearum* (*Fg*) is a major causative agent of FHB of wheat and barley. Wet and warm weather conditions during anthesis favour disease development in wheat. Once *Fg* land on wheat heads, they germinate and penetrate the spikelets through openings between lemma and palea. The pathogen subsequently colonises the head by moving from one spikelet to another. In addition to being responsible for substantial yield losses worldwide each year, mycotoxins produced by cereal infecting fusarium pathogens are harmful to humans and animals if contaminated food and feed is consumed (Goswami & Kistler, 2005).

Understanding cellular processes and identification of genetic elements involved in wheat infection may help in the development of new methods for disease control. Transcriptome profiling has been widely used to interrogate *Fg* transcriptome during infections of wheat heads (Lysøe *et al.*, 2011; Harris *et al.*, 2016; Brown *et al.*, 2017), and stems (Stephens *et al.*, 2008; Zhang *et al.*, 2012) as well as barley (Lysøe *et al.*, 2011; Harris *et al.*, 2016). In fact, the interaction between cereals and *Fg* is one of the most widely studied transcriptome analysis of any plant-microbe interactions (Kazan & Gardiner, 2018). However, transcriptome analyses alone cannot always accurately explain molecular changes as there is not always a clear relationship between transcript and protein abundances (Kage *et al.*, 2020). Recently, a technique called ribosome profiling (ribo-seq), which allows the study of translation on a scale comparable to traditional transcriptomics approaches, has been developed (Ingolia *et al.*, 2009). In this method, ribosomes on mRNAs are immobilised during translation and samples are treated with nucleases to obtain ribosome protected RNA fragments (RPFs). Deep sequencing of RPFs gives quantity and positional information of the ribosomes on transcripts (Kage *et al.*, 2020). Three nucleotide periodicity (Three nucleotide stepwise movement of ribosome on RNA) feature of RPFs helps in identifying unannotated translational events (Liu *et al.*, 2013; Lei *et al.*, 2015; Hsu *et al.*, 2016; Wu *et al.*, 2019) such as translating upstream ORFs (uORFs), downstream ORFs (dORFs), and other ORFs within annotated coding or annotated non-coding regions of the genome. These novel translating ORFs are often missed from genome annotations due to the assumptions underlying most analysis methods (e.g. only the longest ORF within a transcript is most likely to be translated) (Basrai *et al.*, 1997; Claverie, 1997). Hence, ribo-seq can be used for genome-wide experimental identification of actively translating ORFs in an unbiased manner (Wu *et al.*, 2019).

Transcriptomic studies of plant pathogens during various infection stages on host plants have revealed many new insights into potential virulence mechanisms employed by pathogens. However, if transcriptional reprogramming during plant-microbe interactions is correlated at the translational level remains largely unexplored (Kage *et al*., 2020). To answer this question, we performed a genome-wide translatome analysis of *Fg* during wheat head infection using ribo-seq. We discovered that translational responses in *Fg* are moderately coordinated with transcriptional responses during infection of wheat heads. We identified and characterised the function of a translationally regulated *Fg* gene in pathogen virulence. Also, we identified numerous unannotated translation events and examples of regulation at the translational level. Therefore, the experimental strategy used here should be widely applicable to study global translation reprogramming in other plant pathogens during host invasion.

## Materials and Methods

### Culturing *F. graminearum* and sample preparation for ribosome profiling

The *Fg* isolate CS3005 was grown on mung bean agar plate for 1 week by placing a small PDA plug from a water culture stock onto the plate. A small plug from one-week old plate was then transferred to a fresh mung bean agar plate and allowed to grow for 12 days. Two repeated experiments were conducted, and 5 plates were used for each repeat. Fungal samples were collected by scraping the mycelium from the plate using glass slides and immediately immersed in liquid nitrogen and ground into fine powder and stored at −80◻C for future use.

Approximately 0.3 g of fungal tissue powder was resuspended in 1.5 ml of polysome extraction buffer (PEB)/lysis buffer [50 mM Tris-HCl (pH 8), 0.2 M Sucrose, 0.2 M KCl, 15 mM MgCl_2_, 2% (v/v) Polyoxyethylene (10) tridecyl ether (Sigma P2393), 1% triton X-100, 100 μg/mL chloramphenicol, 20 mM 2-Mercaptoethanol, 100 μg/mL cycloheximide, and 10 unit/mL DNase I] and incubated on ice for 10 min with gentle shaking. The lysate was centrifuged at 15,000 × g for 12 min and the supernatant was transferred to new tubes by dividing into 1 ml and 0.5 ml aliquots, flash frozen and stored at −80◻C.

### Plant growth, pathogen inoculation and sample preparation

Four seeds of wheat cultivar Kennedy were sown in each pot in a controlled environment facility (CEF) at CSIRO, St Lucia, Australia. The experiment was laid out in a randomized complete block design (RCBD) with three biological replications over time with twelve plants in three pots as experimental units. Plants were regularly watered, and the room was maintained at 25/18 °C (±1°C) day/night temperature and 65/80% (±5%) day/night relative humidity with a 14-h photoperiod.

For pathogen inoculations, macroconidia from the *Fg* isolate CS3005 grown on mung bean agar plates as described above were harvested and the spore count was adjusted to 1◻×◻10^5^ macroconidia ml^−1^ using an automated cell counter (Cellometer Auto 2000, Thermo Fischer). At 50% anthesis stage, 15 spikes were selected from each replicate and three alternative spikelets were inoculated with 10 μL of macroconidia or mock solution (distilled water + Tween20). Tween20 was added (0.1% v/v) to the spore suspension prior to use. Inoculated spikes were covered with water sprayed zip-lock bags for maintaining high humidity and bags were removed 48 h post inoculation (hpi). At 3 days post-inoculation (dpi), pathogen inoculated spikes were carefully harvested and immediately placed on ice and brought into the laboratory. The region with inoculated and uninoculated spikelets was cut separated and the rachis and the spikelet material were immediately frozen in liquid nitrogen separately and stored at −80◻C until further use. Spikelet samples were finely ground using a pre-chilled pestle and mortal in liquid nitrogen and 0.3 g of powdered sample was ground further in 1.5 ml PEB using an autoclaved and pre-chilled small pestle and mortal until the material was completely dissolved. Remaining sample preparation steps for ribosome profiling were the same as fungal sample preparations explained above.

### Isolation of ribosome protected footprints and ribo-seq

For footprint purification, 1 ml of stored lysate was divided into 0.5 ml aliquots and 6.5 μL of RNase I (100U/μL) was added to each tube (Thermo Fisher Scientific; AM2294). The digestion was carried out at room temperature for 1 h with gentle rotation and stopped by adding 15 μL SUPERase·In RNase Inhibitor (Thermo Fisher Scientific; AM2696). The digestion mixture was layered onto 0.4 ml of sucrose cushion (containing 40 mM Tris-HCl (pH 8), 1 M sucrose, 0.1 M KCl, 15 mM MgCl_2_, 100 μg/mL chloramphenicol, 10 mM 2-Mercaptoethanol, 100 μg/mL cycloheximide, 20 U/ml SUPERase·In) in thickwall 11 x 34 mm polycarbonate tubes (Beckman Coulter Life Sciences; 343778). Samples were weighed and balanced by adding PEB and centrifuged in TLA100.3 rotor at 100,000 rpm for 1.4 h in a pre-chilled ultracentrifuge (Beckman coulter Optima MAX-XP Ultracentrifuge). The supernatant was decanted, and the pellet washed with nuclease free water and resuspended in 0.2 ml nuclease free water containing 10% SDS. RNA was extracted from the resuspended pellet by adding 1 ml Triazol reagent (Invitrogen), vortexed and kept rotating in room temperature for 10 min and 0.2 ml chloroform was added, vortexed and centrifuged at 13,000 ×g for 20 min at room temperature. The aqueous upper phase was transferred to a new tube and 0.7 ml isopropanol was added to precipitate the RNA for overnight at −20◻C. Tubes were centrifuged at 15,000 ×g for 45 min at 4 ◻C to pellet the RNA. The pellet was resuspended in 7 μL of 10 mM Tris buffer (pH 8) and stored at −80◻C.

Footprints were purified by running RNA samples on a 15% TBE urea PAGE gel. To each 7 μL RNA sample, 5 μL 2× denaturing sample loading buffer was added and prepared ladder sample for two lanes. Samples were denatured for 90 s at 80°C, then loaded onto a polyacrylamide gel with size ladders on either side of the RNA samples and separated by electrophoresis for 65 min at 200 V. Gels were stained for 10 min with 1× SYBR Gold in 1× TBE running buffer (5 μL of 10,000× stain/50 ml of TBE buffer) on a gentle shaker and the bands between ~20-35 nucleotides were excised and transferred to a low-bind RNase-free microfuge tube. RNA was extracted from gel slices by adding 400 μL gel extraction buffer (300 mM NaOAc pH 5.5, 1mM EDTA and 0.25% (v/v) SDS), freezing the samples on dry ice for 30 min and thawing at room temperature overnight with slow rotation on a nutator mixer. The extracted RNA was precipitated by adding 1.5 μL GlycoBlue and 500 μL isopropanol at 20°C overnight (McGlincy & Ingolia, 2017). The size-selected RNA was resuspended in 15 μL 10 mM Tris pH 8 and transfer to a RNase-free microfuge tube and stored at −80°C. Next the RNA was subjected to T4 polynucleotide kinase (T4 PNK) treatment to 3‵ end dephosphorylation and 5‵ end phosphorylation, respectively. This reaction was set up in 50 μL (5 μL1x T4 PNK Buffer, 1 μL 20 U SUPERase·In, 1 μL 10 U T4 PNK and 43 μL of diluted RNA) at 37 ◻C for 1 h and thereafter heat inactivation of the enzyme was achieved by incubating for 10 min at 70 ◻C. RNA was precipitated using sodium acetate and isopropanol. NEB’s NEBNext® Multiplex Small RNA Library Prep Set was used for preparing small RNA libraries which were normalized to 2 nM and pooled for sequencing on Illumina NovaSeq 6000 with 100 bp single end reads.

### RNA isolation, processing and RNA-Seq

For RNA isolation, 1 ml Trizol was added to 0.5 ml stored lysate, and RNA was purified as explained above with final resuspension in 20 μL 10 mM Tris buffer (pH 8) and stored at −80 ◻C. RNA samples were treated with DNase I (1 U/μL, Thermo Fisher Scientific; EN0521) by incubating at 37◻C for 30 min and ethanol precipitation was performed. Approximately 8-10 μg of DNA free total RNA (20 μL) was randomly fragmented with equal an volume of 2× alkaline fragmentation solution (2 mM EDTA, 10 mM Na_2_CO_3_, 90 mM NaHCO_3_, pH ≈ 9.3) and the reaction was incubated at 95 °C for 25 min. Fragmentation was stopped by the addition of 0.56 ml ice-cold precipitation solution (300 mM NaOAc pH 5.5, plus GlycoBlue (Ambion) as a co-precipitant), followed by the addition of 0.6 ml of isopropanol and precipitated on ice for 1 h RNA was then pelleted by centrifugation for 30 min at 20,000× g, 4°C in a tabletop centrifuge. The supernatant was removed by pipetting and the pellet air-dried for 10 min before resuspension in 15 μL of 10 mM Tris pH 8. The fragmented RNA was subjected to T4 polynucleotide kinase (T4 PNK) treatment to 3‵ end dephosphorylation and 5‵ end phosphorylation as stated above. RNA was purified using RNA Clean & Concentrator kit (Zymo Research; R101750) followed by phenol-chloroform method detailed in the kit protocol. NEB’s NEBNext® Multiplex Small RNA Library Prep Set was used for preparing small RNA libraries and libraries were normalized to 2 nM and pooled for sequencing on the Illumina NovaSeq 6000 with 100 bp single reads.

### Data analysis

Detailed method is given in Supporting information Method S1.

### Evolutionary analysis of sORFs, and uORF analysis and microRNA target prediction

Detailed method is given in Supporting information Method S2.

### Mutant development, growth determination, virulence assays and ribo-seq analysis of *Fg dicer2*

Detailed method is given in Supporting information Method S2 and S3.

## Results

### Assessment of ribo-seq and RNA-seq data quality

Here, we used RNA-seq and ribo-seq to study transcriptional and translational changes in *Fg* during FHB by comparing non-infectious growth in axenic culture (control) with infectious growth (*in planta*) at 3 dpi (Fig. **1a**). Data quality was analysed using Ribo-seQC (Calviello *et al.*, 2015) and featureCounts (Liao *et al.*, 2014). In total, 33% of the ribosome protected footprint (RPF) and 18% of RNA reads representing 5-12 million reads were uniquely mapped to the *Fg* genome of *in planta* samples (King *et al.*, 2015) per library (Fig. **1b**, Table **S1**). The majority of mapped nuclear RPFs, which were 32 nt or 33 nt-long (Fig. **1c**), were slightly longer than previously reported for nuclear RPFs in yeast, *Arabidopsis* and tomato (28 nt) (Ingolia *et al.*, 2009; Hsu *et al.*, 2016; Wu *et al.*, 2019). Around 78% of the RPF reads were mapped to coding regions (CDS) of the most recently annotated version of the *Fg* PH1 genome (King *et al.*, 2017), with considerably fewer reads mapping to UTRs (6.6%) and unannotated regions (15.6%) of the genome (Fig. **1d**). As expected, RPFs mostly distributed within CDS showed strong three-base periodicity, a feature of translating ribosomes (Fig. **1e**). Pearson correlation analysis showed that biological replicates within control and infection conditions were tightly correlated for both ribo-seq and RNA-seq data (r>0.91, Fig. **S1**). Similarly, ribo-seq and RNA-seq data within control and infection conditions showed significantly high positive correlation (r = 0.78 and r = 0.81, respectively) (Fig. **1f**). The *Fgdicer2* mutant ribo-seq data quality is presented in Fig. **S2**.

**Figure 1:**
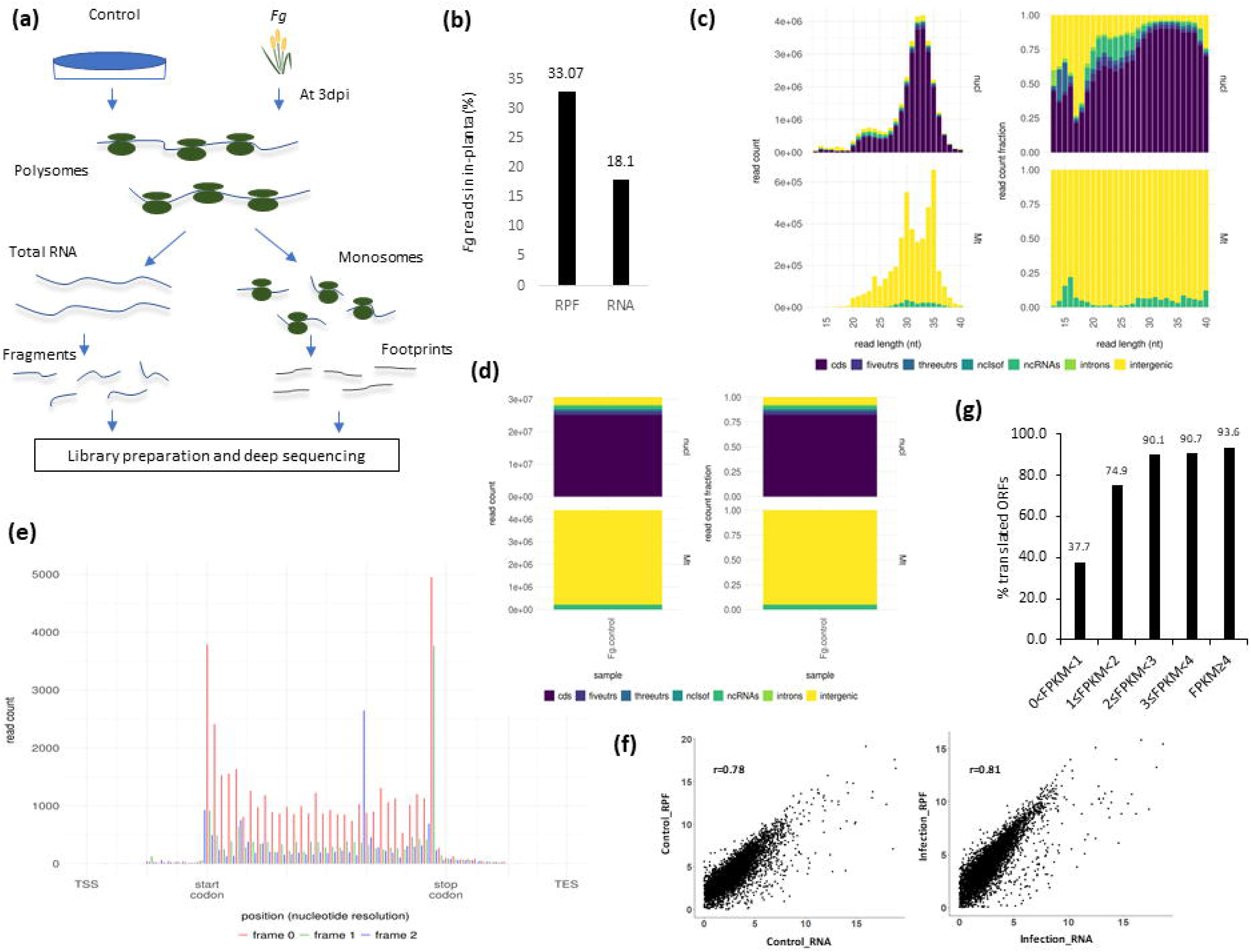
Experimental workflow and data QC. **(a)** Overview of the RNA-seq and ribosome profiling experiment. **(b)** The proportion of *Fg* ribosome protected footprint (RFP) and RNA reads originated from infected wheat heads. (**c)** RPF read length distributions into different genomic locations in *Fg*. **(d)** Percent read distributions into different genomic features in *Fg*. **(e)** Meta-gene analysis of 30 nt RPFs near start and stop codon showing three nucleotide periodicity in *Fg*. **(f)** The Pearson-correlation analysis between ribo-seq and RNA-seq data during control (r=0.78) and infection (r=0.81) conditions. **(g)** Translated ORFs from annotated protein-coding transcripts having different levels of expressions. CDS – coding sequences, utr – untranslated region, RNA – transcriptional level, RPF – translational level.

### Highly transcribed genes are more likely to be preferentially translated in *Fg* during wheat infection

Ribo-seq allows the detection of actively translating ORFs from both annotated and unannotated transcripts. Accordingly, we identified actively translating *Fg* ORFs in our dataset using RiboTaper (Calviello *et al.*, 2015). Among 594 annotated protein-coding *Fg* transcripts having detectable but low expression levels (0< FPKM <1 in RNA-seq) during wheat infection, only 224 (37.71%) were detected as being translated (Fig. **1g**). Out of 101 annotated transcripts having medium (2 ≤ FPKM < 3) expression levels, 91 (90.09%) had actively translating regions and of 3483 annotated transcripts with high (≥4 FPKM) expression levels, 3260 (93.59%) transcripts were detected to have translating regions (Fig. **1g**). This finding suggests that highly expressed transcripts are more likely to be preferentially translated. In addition to translating ORFs identified from coding regions, we detected 36 uORFs and 11 dORFs within the annotated 5‵ and 3‵ UTR regions of annotated *Fg* genes during wheat infection respectively (Table **S2**). However, our uORF and dORF counts are likely underestimates because, out of 14160 transcripts predicted to be encoded by the *Fg* (PH1) genome (King *et al.*, 2015), only 6218 and 5299 currently have annotated 5‵ and 3‵ UTRs, respectively. Thus, RiboTaper might have missed some translated ORFs due to their short lengths and/or low expression levels. We also identified 46 ORFs starting upstream from their originally annotated start sites (Fig. **S3a**, Table **S3**). In addition, some ORFs appeared to use start sites downstream from their annotated start sites (Fig. **S3b**).

### Bulk codon usage analysis in *Fg*

Owing to the accurate definition of coding sequences, riboseq data enable the calculation of, precise codon usage preferences. Therefore, as shown in Fig. **2a, b**, we calculated individual and bulk codon usage summed up over all the positions in *Fg*. This analysis revealed two major groups of codons preferred during translation in *Fg*. The first group contained codons ending with T (GGT, GCT, TCT) while the second group contained codons ending with G or C (GTC, ATC, CTC, TTC, TAC, TGC, CCC, ACC, AAC, CAG, CGC, CAC, AAG, GAC, GAG). This analysis showed that 15 out of 18 amino acids (two amino acids Met and Trp only have one possible codon) favour either G or C at the third base in a codon, whereas only 3 favour either A or T and when there is a C in the second base, and G is avoided in the last codon base as CG combination may lead to methylation (Zhou *et al.*, 2016; Redwan *et al.*, 2019).

**Figure 2:**
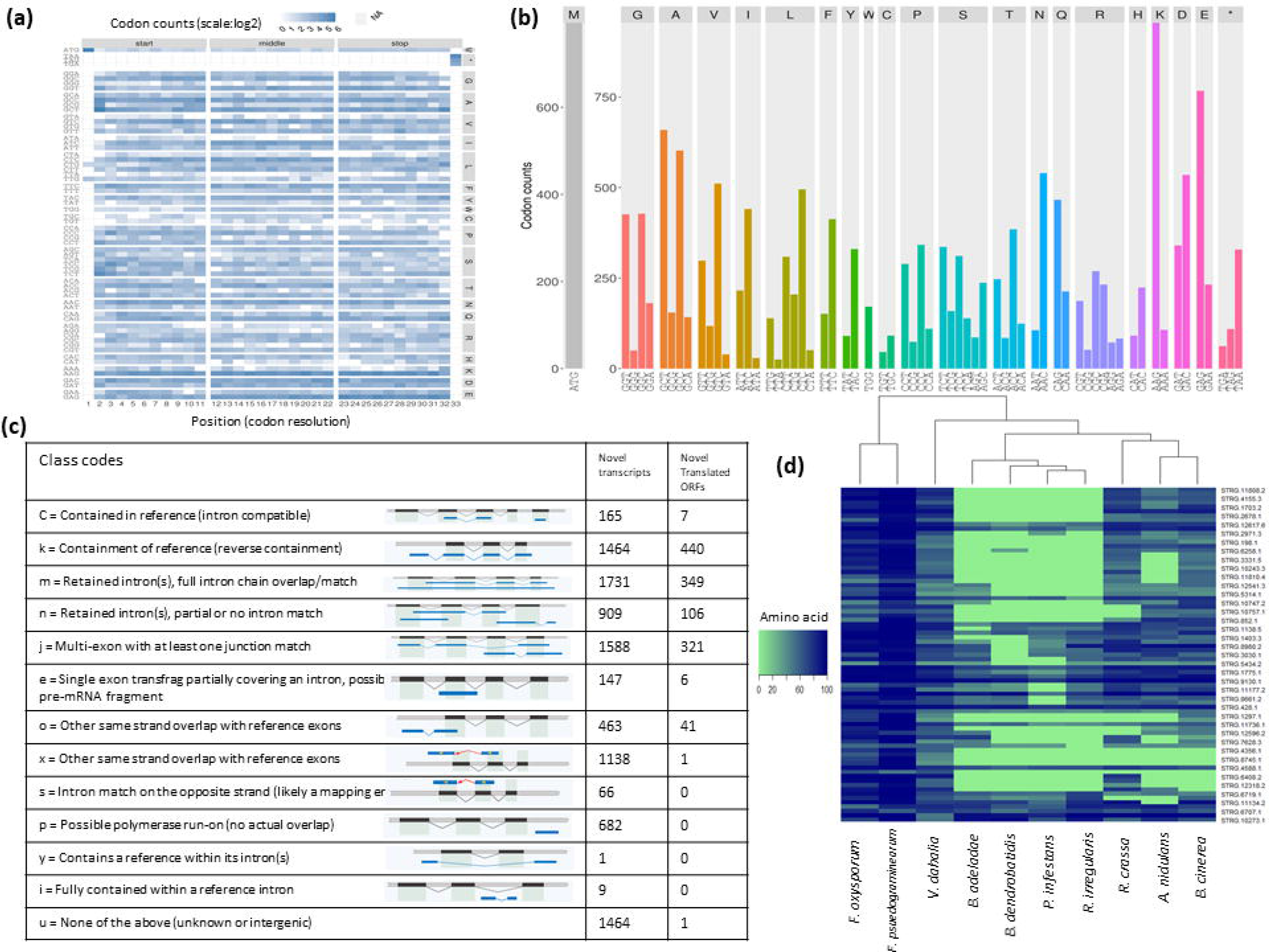
Codon usage and translational landscape in *Fg*. **(a)** Codon usage within CDS of protein coding genes. Position specific codon usage values are calculated for each 11 codons from the first, middle and the last region of CDS. **(b)** Bulk codon usage. The codon usage data were summed up over all CDS positions. Here, codon counts show codon occurrences for each position. **(c)** A summary of novel transcripts and ORFs identified based on Ribotaper analysis in each class of the newly assembled transcripts. Novel transcripts were assembled based on a reference-guided transcriptome assembly using stringtie and gffcompare. The figure was adapted from the gffcompare website (Pertea et al., 2016). **(d)** Evolutionary conservation of small ORFs (sORFs) in *Fg*. Amino acid sequence identities were used to identify homologs in nine diverse different fungal genomes and an oomycete genome. This figure shows only the representative sORFs to make it visible.

### Ribo-seq assisted discovery of novel translating ORFs in *Fg* under control conditions

As genome annotations are highly dependent on automated tools, the availability of RNA-seq and ribo-seq data provides an opportunity to improve genome annotations. Therefore, next, we used our control ribo-seq data to investigate the translational landscape in *Fg* CS3005 to improve genomic resources of this extensively used isolate (Gardiner *et al.*, 2009; Gardiner *et al.*, 2010; Gardiner *et al.*, 2014; Walkowiak *et al.*, 2016; Wang *et al.*, 2017; Beccari *et al.*, 2018). We combined the 100 bp paired-end RNA-seq data generated in our previous study from CS3005 (Accession PRJNA530073) with the data from the present study to develop a reference-guided transcriptome assembly for this isolate. We then compared the new gtf file obtained with the CS3005 annotation (Gardiner *et al.*, 2014) using the gffcompare software (Pertea & Pertea, 2020) to identify previously unannotated transcripts in CS3005. All novel transcripts that potentially encode for new proteins were then combined with the existing annotation of CS3005 to detect actively translating ORFs using RiboTaper. These new proteins were grouped into different classes by gffcompare based on their genomic position and strand information relative to existing gene features (Fig. **2c**). Out of 3791 transcripts with reasonable expression levels (FPKM>1), we detected 2771 (73%) transcripts having translated ORFs. Furthermore, we identified 1272 novel translating ORFs (nORFs) from newly assembled transcripts and grouped them into nine different classes according to gffcompare (Pertea & Pertea, 2020) (Fig. **2c**, Table **S3**). Of these, 157 (12%) were shorter than 100 amino acids (hereafter termed novel sORFs) while the average length of the remaining novel ORFs (88%) was 468 amino acids. Next, we determined functional categories of putative proteins encoded by these novel ORFs. Of these 1272 novel translating ORFs, 166 were predicted to encode putative effectors using EffectorP2 (Sperschneider *et al.*, 2018) (Table **S3**). In addition, 26 encode different classes of putative CAZymes (Carbohydrate-Active enZYmes) predicted based on the dbCAN meta server - automated CAZyme annotation (Zhang *et al.*, 2018) (Table **S3**). Differential expression analysis of nORFs between control and infection conditions showed that 15 of the novel ORFs were accordantly and 3 were discordantly expressed at transcriptional and translational levels in *Fg* during wheat head infection (Table **S3**).

### *Fg* sORFs are evolutionarily conserved

If novel sORFs produce functional proteins, then it is likely that they might be evolutionarily conserved in different fungal organisms. To test this hypothesis, we performed tblastn analyses using 157 sORFs we discovered in *Fg* against 10 diverse fungal and oomycete genomes (Fig. **2d**; Table **S1**). We found that out of 157 sORFs, 144 had at least one homolog in other taxa; some of them are highly conserved in all the genomes (Fig. **2d**). Notably, conserved patterns among the homologs, which largely reflect their phylogenetic relationship, indicate they are unlikely to be false positives. These findings show that the unannotated sORFs identified in this study likely encode functionally important proteins.

### Common and contrasting features of *Fg* transcriptome and translatome during FHB

Differential gene expression analysis using cuffdiff (Trapnell *et al.*, 2012) revealed 1968 significantly differentially expressed genes (DEGs) at the RNA level between control (non-infectious growth) and *in-planta* (infection or infectious growth) conditions. Of these, 716 were upregulated while 1252 were downregulated during infection (Fig. **3a**, Table **S4**). We also identified 3129 differentially translated transcripts (DTTs) during infection by comparing ribo-seq datasets from control and *in-planta* conditions. Of these 3129 genes, 1388 were upregulated, and 1741 genes were downregulated during infection (Fig. **3a**, Table **S4**). The numbers of transcriptionally and translationally downregulated genes were higher than upregulated genes, indicating a global reduction in gene expression during infection at both levels. Further 35.5% of upregulated DEGs overlapped with upregulated DTTs, whereas 43.8% of DEGs overlapped with downregulated DTTs (Fig. **3b**).

**Figure 3:**
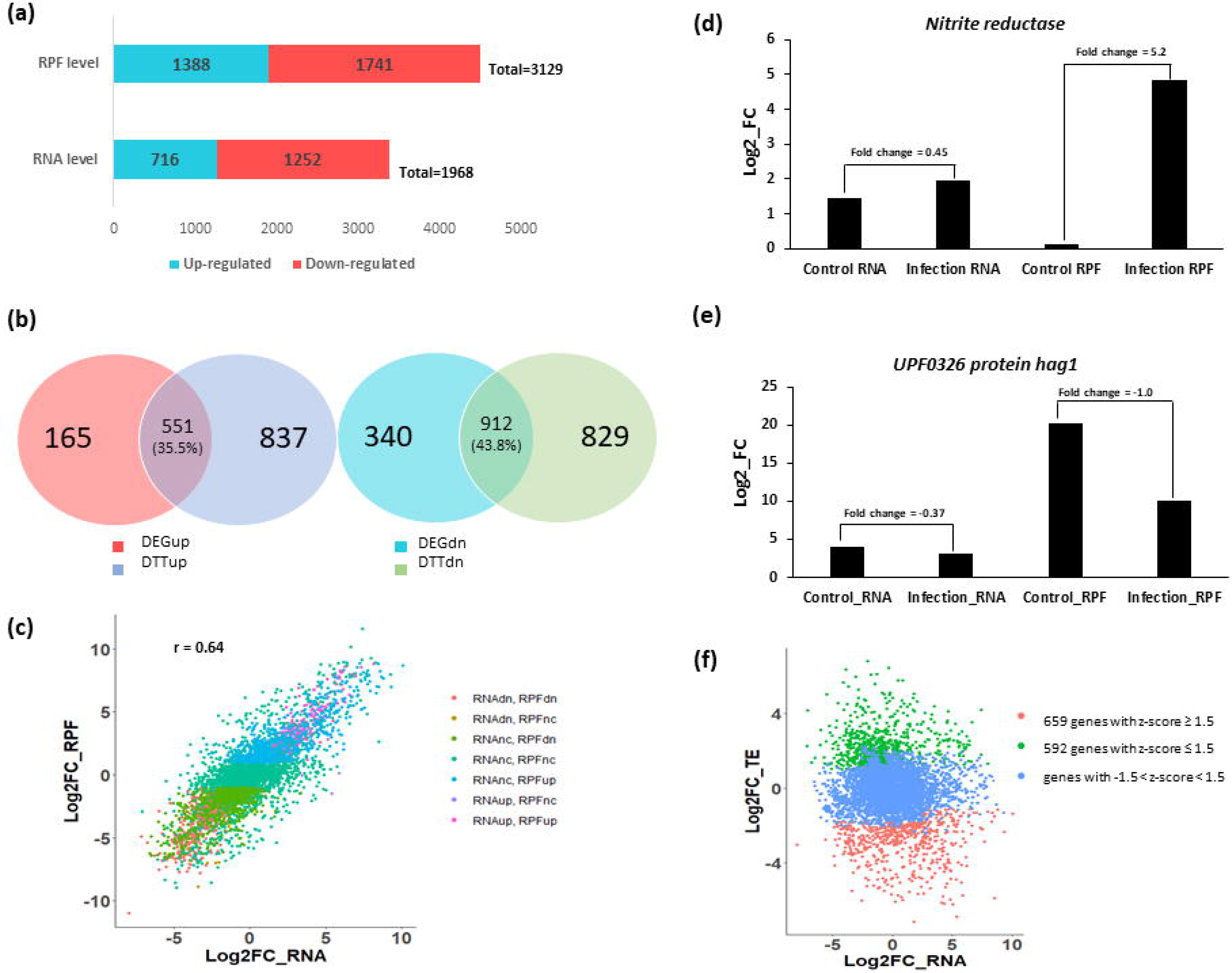
Transcriptional and translational changes during FHB in *Fg*. **(a)** Number of differentially expressed genes (DEGs) at transcriptional and translational levels during FHB in *Fg*. **(b)** Relationships between DEGs at transcriptional and translational levels. The data presented here derived from Figure 2a. **(c)** The relationship between RNA and RPF fold changes**. (d-e)** Discordantly regulated genes at transcriptional and translational levels, translationally upregulated **(d)** and translationally downregulated genes **(e)** without significant changes at transcriptional levels. **(f)** The relationship between RNA and TE fold changes. RNA – transcription level, RPF – translational level, TE – translational efficiency, up – upregulated, dn – downregulated, nc – no change.

Gene ontology (GO) term enrichment analysis (FDR<0.05) showed that the genes transcriptionally and translationally upregulated during infection compared to controls were commonly enriched for processes such as oxidation-reduction, carbohydrate and small molecule metabolism, cellular response to chemical stimuli, organic substance catabolism, cellular oxidant detoxification, and cellular response to oxidative stress (Table **S5**). Among the genes transcriptionally and translationally downregulated, functional categories such as regulation of cellular and biological processes, regulation of transcription, regulation of gene expression, heterocycle biosynthetic process, aromatic compound biosynthetic process, regulation of primary metabolic process, cellular macromolecule metabolic process and RNA biosynthetic processes were enriched (Table **4**).

### Translational reprogramming in *Fg* during FHB

Next, we investigated how transcriptional and translational regulation of gene expression is correlated during infectious growth of *Fg* in wheat heads. These analyses showed that transcriptional changes were moderately correlated (r = 0.64) with translational changes (Fig. **3c**). Two examples of exceptions to this correlation were *Nitrite reductase (FgNiR)* (Fig. **3d**), which was translationally but not transcriptionally upregulated, and *UPF0326 protein hag1* (Fig. **3e**), which was translationally downregulated without significant corresponding transcriptional changes. We then classified the genes regulated at transcriptional and/or translational levels (6788 genes) into seven different possible combinations of these two measures (Fig. **3c**, Table **S6**). Of these, 5257 (77.5%) genes showed correlated expression between transcription and translation (i.e., either transcriptionally and translationally upregulated, downregulated or did not change) (Fig. **3c**, Table **S6**). Discordant transcriptional and translational regulation (transcriptionally up and translationally no change, transcriptionally no change and translationally up, transcriptionally down and translationally no change and transcriptionally no change and translationally downregulated) was observed for 1531 (22.5%) genes (Fig. **3c**, Table **S6**). Interestingly, most trichothecene toxin cluster genes (TRI1, TRI4, TRI6, TRI8, TRI10, TRI11, TRI12, and TRI14) are concordantly translationally and transcriptionally upregulated with the exception of TRI6 (unchanged at both levels) and TRI9, which showed no change at transcriptional levels but translationally up regulated during wheat infection.

To identify translationally regulated genes during host infection, we calculated TE or the rate of mRNA translation into proteins by normalizing ribo-seq FPKM to RNA-seq FPKM as previously reported (Ingolia *et al.*, 2009). This analysis identified 659 and 592 mRNAs showing increased and decreased TE (z| ≥ 1.5), respectively, across all the samples (Fig. **3f**). We removed the genes with little changes at RNA or RPF levels. This refined the absolute numbers to 121 genes with increased TE (TEup) and 195 genes with decreased TE (TEdn) during infection (Table **S6**). GO analysis (p<0.05) of these genes showed that genes with TEup were enriched for polysaccharide metabolic processes, mitochondrial translation, mRNA splicing and cyanide catabolic processes, whereas TEdn genes were enriched for oxidation-reduction, secondary metabolic processes, cellular copper homeostasis and siderophore metabolic processes (Table **S6**).

### Nitrite reductase, a gene with high translational efficiency, is important for virulence in *Fg*

The identification of *FgNiR* as being translationally regulated during infection led us to hypothesise that this gene might be important for infection. To test this possibility, we created two independent mutants of *FgNiR* using the split marker method (Fig. **S4**). To examine the function of *FgNiR* in utilization of different sources of nitrogen, the mutants were grown on minimal media containing nitrite, nitrate, glutamine, urea, or ammonium. At four dpi, both mutants exhibited comparable growth on all the media except on nitrite and nitrate containing media, suggesting these mutants were deficient in using nitrate and nitrite as nitrogen sources due to the disruption of *FgNiR* (Fig. **4a**). Furthermore, growth rates of the mutants were determined by growing them in liquid minimal media supplemented with glutamine as nitrogen source. These experiments revealed there was no difference up to 48 h but differences were observed between 48 and 72 h in the growth patterns of the mutants compared to the wildtype *Fg* (Fig. **4b**). Subsequently, to test the role *FgNiR* in virulence on wheat, a seedling infection assay was performed. *FgNiR* mutants were significantly less virulent than wildtype *Fg* as indicated by reduced shoot length in seedlings inoculated with wildtype *Fg* compared to those mock-inoculated or inoculated with *FgNiR* mutants (Fig. **4c, d**). Fusarium crown rot assay using a seedling infection assay (Gardiner *et al*., 2012) revealed that 32% plants survived in *FgNiR* mutant inoculations whereas none of the plants inoculated with the wildtype strain survived at 13 dpi (Fig. **4e, f**). Furthermore, after inoculation of two central spikelets in a head blight assay, the plants inoculated with either mutant showed significantly reduced symptoms and the spread from the inoculation point to other spikelets (Fig. **4g, h**). Together, these results suggest that FgNiR is an essential component of virulence in *Fg*.

**Figure 4:**
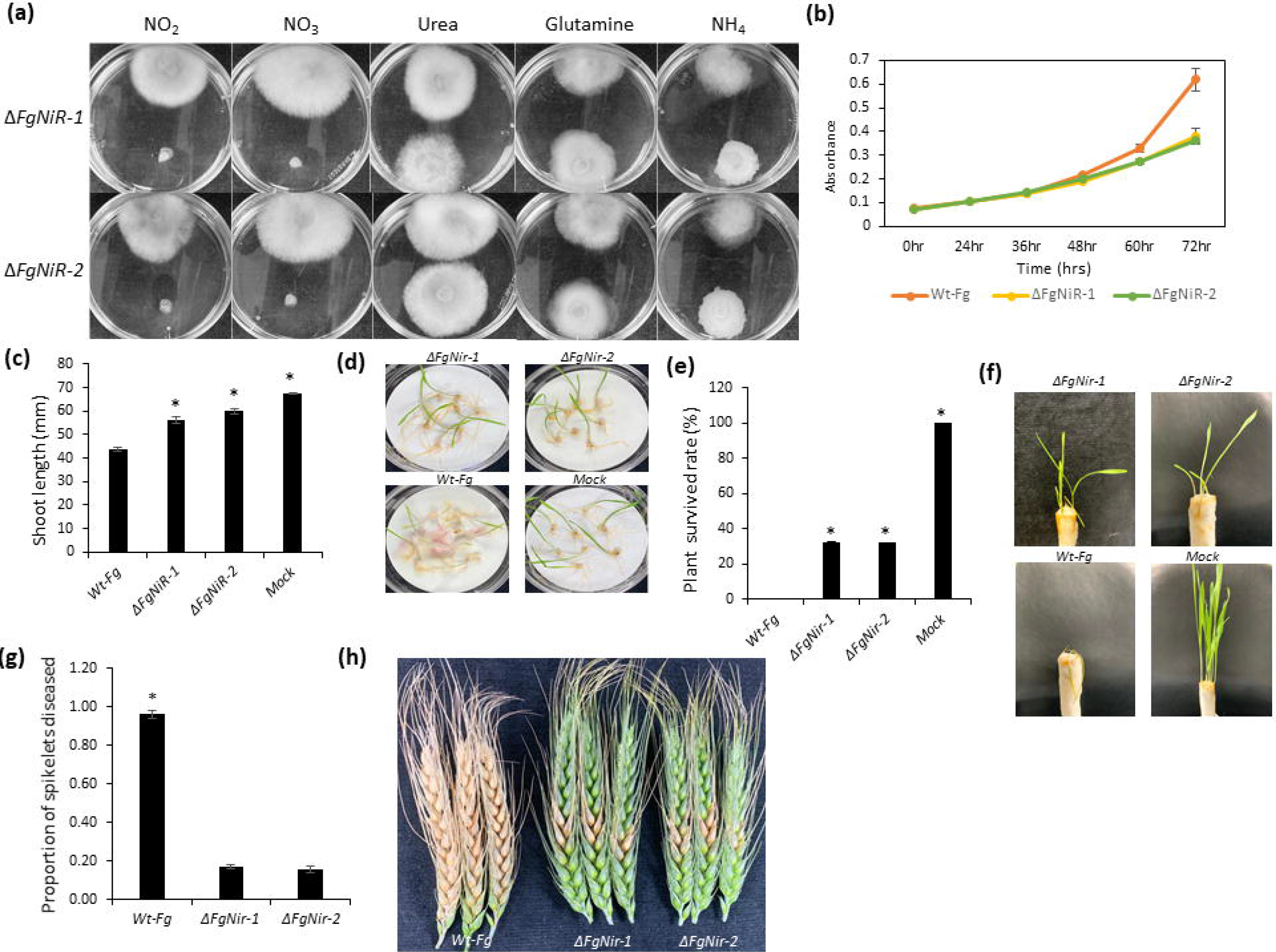
Growth analysis of wildtype *Fg* (CS3005) and *FgNiR* mutants on various nitrogen sources and virulence assays of the mutants towards wheat cultivar Kennedy. In fig **(a)**, the growth of wildtype *Fg* and the *FgNiR* mutants. In both panels of plates, wildtype *Fg* was grown in the upper part while the mutants were grown in the lower parts of each plate. **(b)** Growth of two independent *FgNiR* mutants and wildtype *Fg* in defined media. Error bars represent standard error of the mean for three biological replicates. **(c-d)** Virulence assay of two independent *FgNiR* mutants towards root rot assay at 6 dpi. N = 3 with each replicate consisting of 8-10 plants. **(e-f)** Virulence assay of two independent *FgNiR* mutants towards seedling infection assay. N = 3 with each replicate consisting of 8-10 plants. **(g-h)** Pathogenicity assays of the two mutants of *FgNiR* towards wheat heads. This experiment was replicated three times with 10 spikes each replicate and inoculated pair of middle most spikelets. Photographs show representative infection data. Error bars represent standard error of the mean for three biological replicates and * indicates significance at p<0.05 in pair-wise t-tests.

### Upstream ORFs reduce translation efficiency in *Fg*

The uORFs are known to globally repress translation of their downstream main ORFs (mORFs) in yeast, human, zebrafish, mouse and plants (Ingolia *et al.*, 2009; Liu *et al.*, 2013; Lei *et al.*, 2015; Chew *et al.*, 2016; Spealman *et al.*, 2018). However, how uORFs affect translation in plant pathogenic fungi has not been investigated before. Consistent with previous reports in other organisms, translating uORFs in *Fg* significantly (p=0.01119) reduced TEs of genes associated with uORFs (35 genes in total) as compared to those without uORFs (844 genes in total) during host infection (Fig. **5a**) and a similar trend was observed in control conditions (Fig. **S5**). For instance, FGRAMPH1_01T00769 encoding a hypothetical protein with a translating uORF had a steady transcript level but its TE was significantly reduced during infection most likely due to the presence of an actively translated uORF (Fig. **5b**). To determine whether the genes involved in certain physiological processes are particularly affected by uORFs, we performed a GO enrichment analysis for the genes containing uORFs. This analysis revealed that GO terms such as cellular processes, cellular macromolecule metabolic process, protein metabolic process, cellular protein modification, import across plasma membrane, phosphatidylinositol metabolic process, and GDP binding were enriched among such genes during head infection (Fig. **5c**).

**Figure 5:**
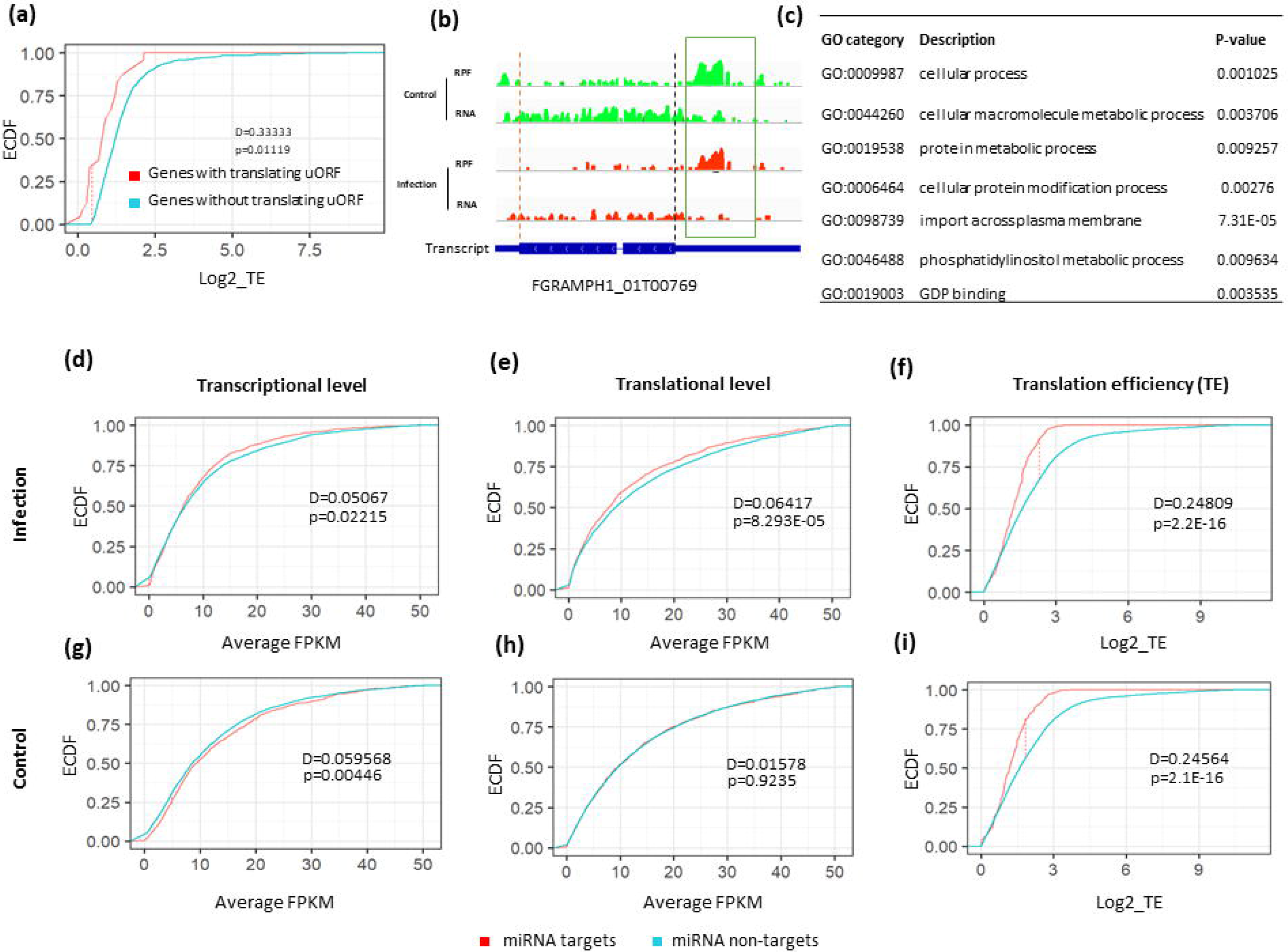
Effect of uORFs and miRNAs on gene expression in *Fg* under control and infection conditions in the form of global distribution of gene expressions. **(a)** Inhibitory effects of upstream ORFs (uORFs) on translational efficiency of downstream main ORFs (mORFs) during infection. Genes (Log2_TE fold change > 0) with and without uORFs were compared here. **(b)** The presence of a translating uORFs found in the upstream region has reduced the translation of the gene FGRAMPH1_01T00769 in *Fg* during infection. Black and red dashed lines indicate annotated start and stop sites, respectively, and green box indicates translating uORF. **(c)** Gene ontology (GO) analysis of uORF containing genes. **(d-f)** Gene regulation by microRNAs at transcriptional level **(d)**, translational level **(e)**, and at TE level **(f)** during infection condition, whereas **(g-i)** depict changes under control conditions. For x-axis in d, e, g and h, only the range from 0 to 50 FPKM and for f and i TE > 0 are shown. ECDF is the empirical cumulative distribution function of the samples. Red dotted line is absolute difference (D) between the two distributions called as K-S statistic. The p and D-values were determined with two-sample Kolmogorov-Smirnov test.

### miRNAs act in *Fg* to control translational efficiency

Post-transcriptional gene expression in eukaryotes is also regulated by miRNAs through mRNA cleavage or stalling the initiation or extension of translation by forming a duplex with translating mRNAs (Yu *et al.*, 2017; Li *et al.*, 2018). In *Fg*, potential roles of miRNAs are not well known. To globally investigate the effect of miRNAs on gene expression at transcriptional and translational level, we acquired 49 previously reported *Fg* miRNA sequences by (Chen *et al.*, 2015) and predicted their targets (Table **S5**) using psRNATarget (Dai *et al.*, 2018). These authors also showed that these miRNAs could regulate the expression of target genes in *Fg*. Therefore, we compared transcriptional and translational abundances as well as TEs of miRNA target and non-target genes. This analysis showed that transcript levels of miRNA targets were significantly (p=0.02215) reduced compared to non-targets during infection (Fig. **5d**). Furthermore, RPF levels (p=8.293×10^−05^) and TEs of miRNA target genes (p=2.2×10^−16^) were significantly reduced as compared to non-targets during infection (Fig. **5e, f**). In contrast, only transcriptional levels (p=0.00446) and TEs (p=2.1×10^−16^) but not translational levels (p=0.9235) of the miRNA targets were significantly reduced compared to non-miRNA targets in control samples (Fig. **5g, h, i**). Thus, we conclude that miRNAs regulate global gene expression at both transcriptional and translational levels to control TE in *Fg* during head infections whereas during control conditions miRNAs only affect transcript levels to modulate TEs of the genes.

### Dicer2 dependent translational regulation in *Fg*

miRNAs are biosynthesised in fungi by different mechanisms using different factors, with dicer2 among such factors involved (Lee *et al.*, 2010). Information on the role of Fgdicer2-dependent miRNA on transcriptional and translational gene expression in plant pathogenic fungi is very limited. Chen *et al.* (2015) reported that the biogenesis of 24 miRNAs, out of 49 miRNAs, was Fgdicer2-dependent. To further investigate the role of Fgdicer2 in miRNA biogenesis, and how the dicer2-dependent miRNAs regulate gene expression at transcriptional and translational level, we created a *Fgdicer2* mutant and performed ribo-seq of *Fgdicer2* and wildtype *Fg* grown in culture. This analysis revealed that there were no changes in the overall distribution of transcriptional gene expression in these genotypes (Fig. **6a**). In contrast, the overall distribution of gene expression at both translational levels and TEs were increased in the *Fgdicer2* mutant compared to wildtype *Fg* (Fig. **6b, c**). This could be due to the loss of Fgdicer2-dependent miRNA-repression. To confirm these results, we pulled Fgdicer2-dependent miRNAs from Chen et al., (2015) and looked at the expression levels of their target genes at transcription, translation, and TE levels. These analyses revealed the same trend observed at genome-wide analysis (Fig. **6d, e, f**). Interestingly, the expression of the trichothecene toxin cluster genes was neither influenced by Fgdicer-dependent nor non-dependent miRNAs in *Fg*.

**Figure 6:**
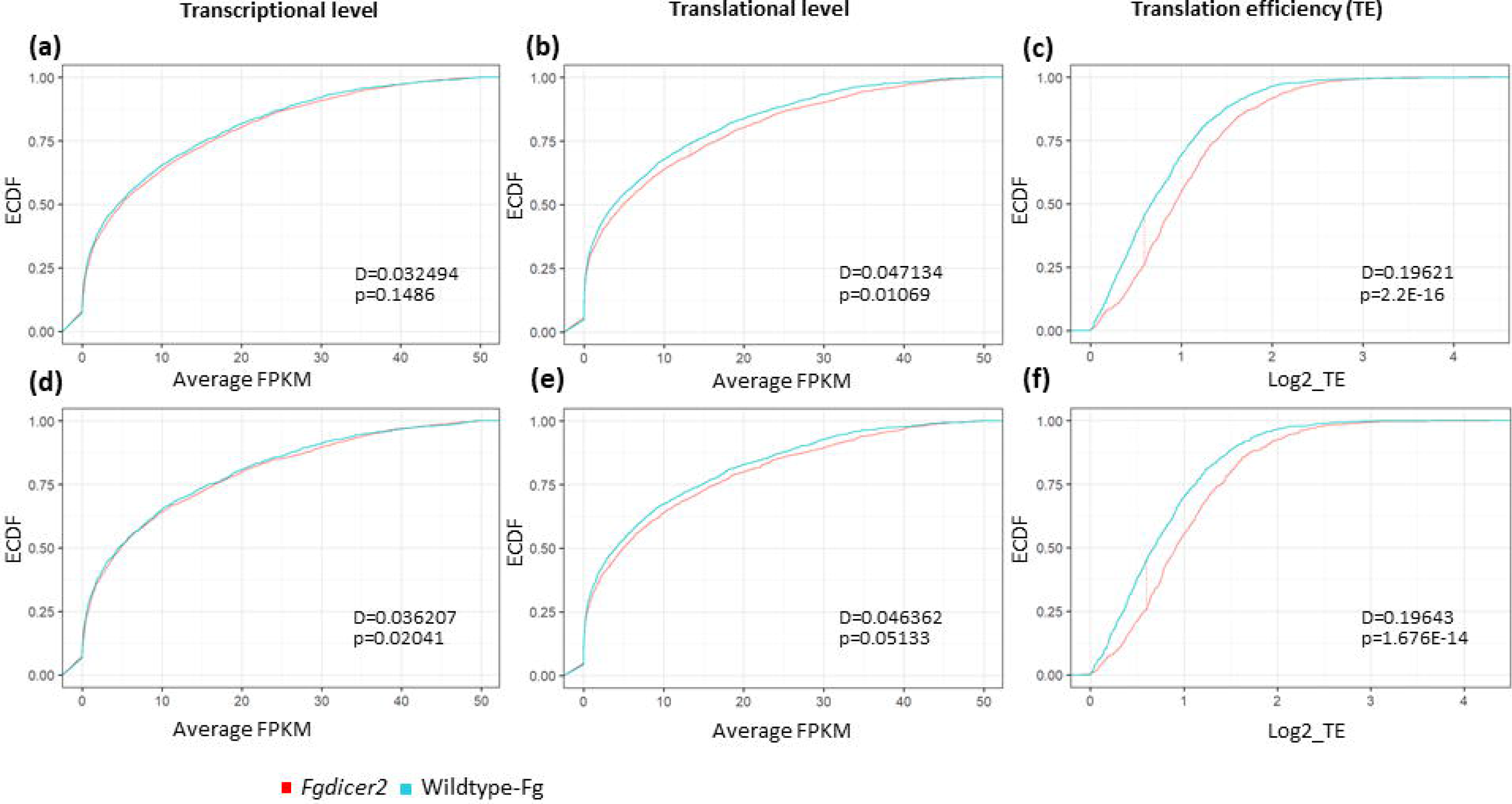
Global gene expression distribution patterns of miRNA targets in wildtype and *Fgdicer2* mutant of *Fg*. **(a-c)** show gene regulation by all the microRNAs at transcriptional level **(a)**, translational level **(b)**, and at TE level **(c)**, whereas (d-f) show changes in gene regulation by Fgdicer2-dependent miRNAs at the transcriptional level **(d)**, translational level **(e)**, and at TE level **(f)**. For x-axis in a, b, d and e, only the range from 0 to 50 FPKM and for c and f TE > 0 are shown. ECDF is the empirical cumulative distribution function of the samples. Red dotted line is the absolute difference (D) between the two distributions called K-S statistic. The p and D-values were determined with two-sample Kolmogorov-Smirnov test.

## Discussion

### Ribosome profiling identifies novel translational events in *Fg*

Our understanding of translational control in fungi is largely limited to yeast. In plant pathogenic fungi, better understanding of translation events associated with pathogenicity is important as it leads to the development of sustainable plant protection strategies. In this study we focussed on *Fg*, an economically important cereal pathogen. To the best of our knowledge, genome-wide analysis of mRNA translation has not been investigated in *Fg*. Therefore, we undertook a combined RNA-seq and ribo-seq analysis to understand gene expression changes at transcriptional and translational levels to understand translational control of gene expression in *Fg*. We found that many translational features are shared between *Fg* and other eukaryotes. For instance, the length of RPFs is around 32-33 nts in *Fg* (Fig. **1c**), which is slightly longer than previously reported RPF length for yeast (28) nt (Ingolia *et al.*, 2009). The hallmark of ribo-seq data, i.e. a 3-nt periodicity (Fig. **1e**), is also observed in *Fg* ribo-seq data (Ingolia *et al.*, 2009; Hsu *et al.*, 2016; Wu *et al.*, 2019). With the help of RiboTaper, we found out that highly expressed transcripts are more likely to be translated during infection in *Fg*. This conclusion is consistent with a previous report in *Arabidopsis* where increased fractions of detectable translated ORFs were found among highly expressed transcripts (Hsu *et al.*, 2016). We were also able to identify new translation events such as uORFs and dORFs which are previously unannotated in *Fg* (Fig. **S2**, Table **S2)**. Additionally, we found novel translated ORFs from a newly assembled transcriptome of isolate CS3005 that encoded proteins such as secreted peptides, CAZymes and effectors (Table **S3**). *Fg* is known to secrete a cocktail of many enzymes, proteins, toxins, and other virulence factors to infect plants (Brown *et al.*, 2017). Secreted peptides and CAZymes target plant cell wall and induces cell death (Brown *et al.*, 2012; Zhang *et al.*, 2012; Sella *et al.*, 2013; Sperschneider *et al.*, 2016) and effectors usually help to evade host defence by suppressing its immune system (Stergiopoulos & de Wit, 2009). Further we discovered many novel sORFs which are highly conserved in fungi and thus likely encode functional proteins. Our observations agree with a report in tomato, where sORFs are highly conserved across different plant genomes, suggesting their functional significance throughout evolution (Wu *et al.*, 2019). Therefore, novel translating proteins identified in this study may potentially act as unique arsenals for *Fg* during the invasion of wheat heads and further studies are required to discover their biological significance. Our study also found that some ORFs starting translation upstream or downstream from their annotated start sites, these observations suggest that not all upstream AUG sites are used as translation initiation sites, consistent with reports in yeast (Ingolia *et al.*, 2009; Smith *et al.*, 2014), tomato (Wu *et al.*, 2019) and *Arabidopsis* (Hsu *et al.*, 2016).

### *Fg* infection of wheat is governed by responses at both transcriptional and translational levels

Our results showed a global reduction in *Fg* gene expression, indicating that transcriptional or translational regulation of gene expression during infection may be due to the repressive effects of miRNAs and/or uORFs (Fig. **5d-e**) to balance between growth- and virulence-associated processes. Furthermore, we found that 22.5% of *Fg* genes are discordantly regulated transcriptionally and translationally during host infection, suggesting that studying transcription alone would not reveal the full extent of gene regulation in this pathogen. GO analyses of transcriptionally and translationally upregulated genes were enriched for carbohydrate and small molecule metabolism and response to oxidative stress, which are involved in energy production to help virulence and fight with oxidative stress caused by plant defense respectively. The regulation of cellular and biological processes was transcriptionally and translationally downregulated. This might be to encounter plant defense aggressively by switching away from growth-related activities. Similarly, GO analysis of only translationally upregulated genes showed that pathways associated with glucan, cellulose and hemicellulose metabolic processes were enriched (Table **S5**). These pathways are important for the pathogen to destroy plant cell walls which act as barriers for pathogen entry. The reason for genes which are regulated only at transcriptional level with no detectable change at translational level is possibly due to the delay between mRNA and protein synthesis because maturation, export, and translation of mRNAs take some time (Liu *et al.*, 2016). In yeast, it has been reported that the transcriptional changes measured at a time point was better correlated with protein abundances at a later time point under sodium chloride stress (Lee *et al.*, 2011), indicating protein expression lags behind transcriptional changes. Other likely explanations for this delay would be due to post-transcriptional regulations (Franks *et al.*, 2017) or the transcript lengths, codon compositions or alterations in translation (Fournier *et al.*, 2010; Gedeon & Bokes, 2012). It is also be possible that fungi modulate transcript levels to reduce translational efficiency to maintain essential house-keeping processes running (Lei *et al.*, 2015) or cells making mRNA pools to use for quick translation when needed (Shenton *et al.*, 2006). Nevertheless, we also detected *Fg* genes exclusively regulated at the translational level with no detectable changes at the transcriptional level. In some cases, increased translation of existing mRNAs contributes to the synthesis of newly required proteins. This process called “translation on demand” can act as an independent translational response, ensuring that proteins are quickly synthesised in response to signals rather than constitutively synthesised (Beyer *et al.*, 2004). Taking all these into consideration, our results show that during infection, *Fg* genes are regulated at transcriptional and/or translational levels either co-ordinately or independently and this may be dependent upon the severity of stress or the extent of necessity.

Furthermore, we identified 121 *Fg* genes with TEup and 195 genes with TEdn during infection (Table **S6)**. *FgNiR* was one of the highly translationally regulated genes and was functionally characterized to show its role in pathogen virulence. Using traditional gene expression analyses alone, this gene could not have been identified as a virulence gene candidate. In the absence of preferred nitrogen sources during infection, fungal pathogens assimilate nitrate or nitrite, the most abundant nitrogen source available in plants, and adapt to nitrogen-limiting conditions through the regulation of secondary nitrogen acquisition (Divon *et al.*, 2006; Song *et al.*, 2007). In *Aspergillus nidulans*, the *NiR* was only expressed under nitrogen starvation and simultaneous presence of nitrite (Marzluf, 1997). Similar observations were made in *F. fujikuroi* where *NiR* was regulated by the transcription factor NirA and the nitrogen-dependent regulator protein AreB (Pfannmüller *et al.*, 2017). Since it is only translationally regulated, it appears that *FgNiR* is expressed in the “translation on demand” mode to cope with nitrogen starvation conditions *in planta* during infection through an unknown mechanism that needs to be further explored. Other translationally regulated genes (TEup or TEdn) could serve as candidates for further investigation of their involvement during wheat infection. Such genes were enriched for mitochondrial translation and mRNA splicing pathways that might lead to the production of novel virulence-related mitochondrial proteins or alternative translation products from mRNA splicing. Overall, our findings demonstrate that TE could be used as a measure in selecting candidate virulence genes for functional characterization.

### Translational regulation of gene expression by uORFs and miRNAs

In our genome wide analysis, we identified translating uORFs that inhibited the translation of downstream mORFs in *Fg* during wheat infection (Fig. **5a, b**). Similar phenomena have been reported in *Arabidopsis* (Liu *et al.*, 2013), maize (Lei *et al.*, 2015), and tomato (Wu *et al.*, 2019). Repression of mORF translation is most likely due to both uORFs and mORFs being on the same transcript and competing for the translation initiation complex. Alternatively, termination of uORF translation hinders the re-initiation of mORF translation (Sachs & Geballe, 2006; Barbosa *et al.*, 2013).

Our study also revealed negative effects of miRNAs on gene expression. *Fg* miRNAs reduced not only transcription, and translation but also TEs of miRNA target genes as compared to non-targets during wheat infection (Fig. **5d, e, f**). miRNAs regulate transcript levels by mRNA decapping or decay and inhibit translation by binding and cleaving transcripts (Bazzini *et al.*, 2012; Hausser *et al.*, 2013). Our observation of transcriptional and translational repression by microRNAs is consistent with previous reports, where miRNAs target transcripts were found to have significantly lower TE in photomorphogenic *Arabidopsis* (Liu *et al.*, 2013). In another study, miRNAs similarly regulated gene expression at both transcriptionally and translationally in tomato (Wu *et al.*, 2019). Several studies have reported the involvement of miRNAs in pathogenesis (Chen *et al.*, 2016; Arora *et al.*, 2017; Wang *et al.*, 2018). If involved in *Fg* pathogenicity, such miRNAs could be potential targets for new fungicides. Interestingly, none of the trichothecene cluster genes were found to be affected by miRNAs in *Fg* and this might be due to the importance of deoxynivalenol for virulence as well as survival of the pathogen in the environment.

Increased gene expression observed in the *FgDicer2* mutants could be due to the loss of Fgdicer2-dependent miRNA-repression, whereas decreased gene expression found in this mutant might be attributed to either inhibitory effects of non-Fgdicer2-dependent miRNAs on gene expression or Fgdicer2-dependent miRNAs positive regulatory effects. Indeed, it was reported that miRNAs can both upregulate and downregulate gene expression (Valinezhad Orang *et al.*, 2014; Huang, 2017). We found that the genes encoding *hypothetical proteins* (FGRAMPH1_01T12183, and FGRAMPH1_01T16367), *polyketide synthase* 3 (FGRAMPH1_01T27727), responsible for the biosynthesis of the perithecial pigment, and *ankyrin repeat protein* (FgANK1 - FGRAMPH1_01T11519), required for host sensing (Ding *et al.*, 2020), were highly upregulated translationally in the *Fgdicer2* mutant. Similarly, the genes such as *polyphosphate synthase* (FGRAMPH1_01T03521), *15-o-acetyltransferase* (FGRAMPH1_01T13105) and *serine/threonine protein kinase* (FGRAMPH1_01T14345) had higher TEs in the *Fgdicer2* mutant compared to the wildtype *Fg*. These observations suggest that biological processes associated with these genes are controlled by Fgdicer2-dependent miRNAs while Fgdicer2-dependent miRNAs do not appear to play a role in controlling the expression of trichothecene biosynthesis genes in *Fg*.

In conclusion, our study provided an in-depth profile of the translational dynamics of *Fg* during wheat infection. As well as characterising known and novel ORFs transcription and translation events during infection, we also discovered possible roles of uORFs and microRNAs in regulating global gene expression in this important pathogen. Further functional analyses of candidate genes identified here might provide novel targets for intervention to reduce Fg-incited crop losses. The knowledge gained here should be applicable into other host-pathogen interactions and may lead to the development of effective pathogen control strategies.

## Acknowledgments

The first author is a recipient of early research career (CERC) postdoctoral fellowship from CSIRO, Australia. We are thankful to Jonathan Powell for his suggestions for bioinformatics analysis and Anca Rusu for developing the *dicer2* mutant. The authors would also like to thank Polly Hsu and Hsin-Yen Larry Wu from Michigan State University, USA, Nicholas Ingolia from the University of California Berkeley USA, and Lorenzo Calviello from the University of California, San Francisco, USA for their invaluable suggestions on ribo-seq experiments and/or bioinformatics analysis.

## Author Contributions

UK, DMG, JS and KK planned and designed the research. UK performed the experiments, JS helped in writing scripts and bioinformatics analysis. UK wrote the first draft of the manuscript and UK, DMG and KK contributed to the final draft of the paper.

## Data Availability

The data reported in this paper is deposited at the NCBI short read archive under the accessions PRJNA729871 and PRJNA729812.

## Supporting information Legends

**Method S1:** Data analysis

**Method S2:** Evolutionary analysis of sORFs, and uORF analysis and microRNA target prediction

**Method S3:** Mutant development, growth determination, and virulence assays.

**Method S4:** *Fgdicer2* mutant development and ribo-seq analysis.

**Supporting information Fig. S1:** Correlation for ribo-seq and RNA-seq data within the same treatment.

**Supporting information Fig. S2:** Ribo-seq data quality for wildtype *Fg* and *Fgdicer2* mutant.

**Supporting information Fig. S3:** Novel ORFs identified from ribo-seq data in *Fg*.

**Supporting information Fig. S4:***FgNir* mutant development using split marker strategy and mutant screening by PCR.

**Supporting information Fig. S5:** Effect of upstream ORFs (uORF) on translational efficiency of mORF under control conditions in *Fg*.

**Supporting information Table S1:** Read mapping statistics of *Fg* and reference genomes from Ensembl used for evolutionary analysis of small ORFs.

**Supporting information Table S2:** Translated ORFs in *Fg* during wheat infection.

**Supporting information Table S3:** Translated ORFs in *Fg* under control condition and functional categories of novel ORFs (nORFs) and their expressions at transcriptional and translational level.

**Supporting information Table S4:** Differential gene expression at transcriptional and translational level in *Fg* during wheat infection.

**Supporting information Table S5:** List of miRNAs, miRNA targets and gene ontology (GO) analysis in *Fg*.

**Supporting information Table S6:** List of translationally regulated genes and GO analysis in *Fg* during wheat infection.

**Supporting information Table S7:** Transcriptional and translational gene expressions in wildtype *Fg* and Fgdicer2 mutant.

## Notes

### Competing Interest Statement

The authors have declared no competing interest.

