## Supporting Information for "Translational control of fungal gene expression during the wheat-*Fusarium graminearum* interaction"

**New Phytologist Supporting Information**

Title: Translational control of the infection process of *Fusarium graminearum* on wheat

acceptance date:

**Supporting information Method S1:** **Data analysis**

### The raw RNA-Seq and ribo-Seq reads were adapter and quality trimmed using trimmomatic v0.36 (Bolger *et al.*, 2014). Sequence read quality check was performed before and after trimming using FastQC (http://www.bioinformatics.babraham.ac.uk/projects/fastqc/). The genome and annotation of *Fg* PH1 [ASM24013 v3](http://www.ebi.ac.uk/ena/data/view/GCA_000240135.3).0 were downloaded from Ensembl (http://fungi.ensembl.org/Fusarium_graminearum_gca_000240135/Info/Index). Trimmed and quality checked reads which did not map (Bowtie2 v2.3.4) to a contamination library (rRNA, snRNA and snoRNA) or wheat genome were mapped to the PH1 sequence (King *et al.*, 2017) using TopHat v2.1.1. Transcriptome and translatome abundances and fold changes were calculated using cufflinks and cuffdiff v2.2.2, respectively (Trapnell *et al.*, 2012). Translational efficiency (TE) was calculated by normalizing ribo-seq FPKM to RNA-seq FPKM (Ingolia *et al.*, 2009).

### For RiboTaper analysis in *Fg*, the mapped RNA-Seq reads from this study and from our previous study (Accession PRJNA530073) were merged (BAM files) and used to perform reference guided *de novo* assembly with stringtie v1.3.4d (Pertea & Pertea, 2020). A newly assembled gtf file was compared with CS3005 v1.0 gtf (downloaded from Ensembl) using gffcompare v0.11.2 (Pertea & Pertea, 2020). The new classes (Pertea & Pertea, 2020) transcripts information was extracted from the gffcompare output and merged with CS3005 v1.0 gtf and the resulting merged gtf was used to map RNA-Seq and ribo-Seq reads and downstream RiboTaper analysis. Two replicates of mapped bam files from RNA-Seq and ribo-Seq were merged using SAMtools v1.9.0 (Li *et al.*, 2009) and used for ORF discovery using RiboTaper v1.3.0 (Calviello *et al.*, 2015).

**Supporting information Method S2:** **Evolutionary analysis of sORFs, and uORF analysis and microRNA target prediction**

For evolutionary analysis of sORFs, sequences shorter than 100 amino acid were used to perform the TBLASTN with an e-value cut off of 0.001 (Camacho *et al.*, 2009) against the different fungal/or oomycetes reference genomes downloaded from Ensembl (Supporting information Table S1). Evolutionary analyses were done according to the method described previously (Wu *et al.*, 2019). Amino acid identity was plotted in Rstudio using heatmap3 (Zhao *et al.*, 2014).

To investigate the effect of uORFs on translation of mORFs, we first identified actively translating uORFs in *Fg* using RiboTaper (Calviello *et al.*, 2015) and then compared the TEs (Log2_TE fold change >0) of genes with and without uORFs. For miRNA analysis, previously reported miRNA sequences obtained from Chen *et al*., (2015) were used against CS3005 v1.0 mRNA sequences to predict miRNA targets in *Fg* using psRNATarget (Dai *et al.*, 2018) with SchemaV2 2017 release.

**Supporting information Method S3: Mutant development, growth determination, and virulence assays**

The *FgNir* was developed by split marker recombination method as described earlier (Wang & Tang, 2018) with [nourseothricin N-acetyl transferase (NAT1) as selectable marker with PEG-mediated protoplast transformation](https://plantmethods.biomedcentral.com/articles/10.1186/s13007-019-0526-5) (Desmond *et al.*, 2008). Transformants were validated for mutants of *FgNir* using primers (Primer1F: CAACAAAATCGATCTTATTGAGAG, Primer1R: GATCCCTTCTCCAACGTCTAC, Primer2F: CTCATGACAGCTAAAACCTAC

TAAGGTA, Primer2R: CAAGCAGCAGATGATAATAATGTCCTC). PCR validated two independent knock-out mutants were grown on minimal media (Correll *et al.*, 1987) with different nitrogen sources at 28 °C as previously described (Pfannmüller *et al.*, 2017) for growth determination on solid media. Similarly, growth rate determination was performed in liquid media according to earlier method (Gardiner *et al.*, 2012). For virulence assays mutants were grown on half strength potato dextrose agar media and spore production was carried out in CMC media (Tuite, 1969) as above described. Fusarium crown rot virulence assay was performed using an earlier method (Gardiner *et al.*, 2012) and for fusarium seedling infection assay germinated seeds were inoculating with 5ml of spore suspension in falcon tube for two min by rolling to coat the seeds and 8-10 seeds were placed in petri plates assembled with two layers of filter papers wetted with 10ml of water. Petri plates were sealed and kept in room temperature with 12 hours of light for 6 days. Assays were scored by measuring the shoot length. Fusarium head blight assay was carried out by inoculating a middle most pair of spikelets of wheat heads during anthesis stage and scoring the proportion of diseased spikelets. This experiment was replicated three times with 10 spikes each replication.

**Supporting information Method S4:** **Fgdicer2 mutant development and ribo-seq analysis**

The *Fgdicer2* mutant was created from transformation of CS3005 (as described above) with a PCR product amplified from a synthesised construct for double cross over recombination. The amplified product contained in order; 1000 bp of sequence directly upstream of the Fgdicer2 (FG05_04408) start codon, a spacer with sequence gatgtccacgaggtctctaagcgcgcgattacgtattccgtacgctgcaggtcgac, a Aspergillus nidulans TrpC promoter driving expression of nourseothricin acetyl transferase corresponding to nucleotides 437-1387 of genbank accession AY631958.2 and 992 bp of sequence corresponding to the Fgdicer2 terminator. Transformants were screen for successful deletion of Fgdicer2 using a triplex PCR assay whereby the mutant was identified by the loss of a wild type 421 bp product and gain of a vector specific band of 523 bp (FG05_04408KOscrnF: AATTTCTGGTCGGCCTCCTG, FG05_04408KOscrnR: CGCCTGTAGCAGAGATCTCG, TrpCR: GCTGATCTGACCAGTTGCCT). The *Fgdicer2* mutant and Fg-wildtype were grown on mung bean agar plates and sample was collected as explained in above section with three repeated experiments with 10-15 plates each. Ribo-seq experiment and data analysis was conducted same as we explained in material and methods.

**Supporting information Table S1: Read mapping statistics of *Fg* and reference genomes from Ensembl used for evolutionary analysis of small ORFs**

| Sample | Total raw trimmed reads | Pure reads | Unique mapped reads | % |
| --- | --- | --- | --- | --- |
| Control_RNA1 | 102,880,949 | 56,404,790 | 8,685,616 | 15.39872057 |
| Control_RNA2 | 41,084,065 | 27,616,794 | 5,788,167 | 20.95886655 |
| Control_RPF1 | 133,925,024 | 81,451,329 | 17,153,054 | 21.05926841 |
| Control_ RPF2 | 131,698,916 | 88,451,966 | 13,826,934 | 15.63213869 |
| *In-planta*_RNA1 | 80,133,327 | 33,136,043 | 5,401,032 | 16.29956842 |
| *In-planta*_RNA2 | 93,848,187 | 36,411,779 | 7,275,933 | 19.98236065 |
| *In-planta*_RNA3 | 79,293,178 | 28,405,816 | 5,256,650 | 18.50554126 |
| *In-planta*_ RPF1 | 115,724,564 | 33,957,523 | 10,103,687 | 29.75389872 |
| *In-planta*_ RPF2 | 113,354,455 | 34,391,260 | 12,174,881 | 35.40109028 |
| *In-planta*_ RPF3 | 121,699,232 | 35,845,332 | 11,610,299 | 32.38998874 |
| **Different fungal/or oomycetes reference genomes** | | | | |
| 1. Aspergillus_nidulans.ASM1142v1.dna.toplevel (Ascomycota) 2. Batrachochytrium_dendrobatidis_ jel423_gca_000149865.BD_JEL423.dna.toplevel (Chytridiomycota) 3. Bifiguratus_adelaidae_gca_ 002261195.ASM226119v1.dna.toplevel (Zygomycota) 4. Botrytis_cinerea.ASM83294v1.dna.toplevel (Ascomycota) 5. Fusarium_oxysporum.FO2.dna.toplevel (Ascomycota) 6. Fusarium_pseudograminearum.GCA_000303195.1. dna.toplevel (Ascomycota) 7. Neurospora_crassa.NC12.dna.toplevel (Ascomycota) 8. Phytophthora_infestans.ASM14294v1. dna.toplevel (Oomycota) 9. Rhizophagus_irregularis_daom_181602_gca_000439145.ASM43914v3. dna.toplevel (Zygomycota) 10. Verticillium_dahliae. ASM15067v2.dna.toplevel (Ascomycota) | | | | |

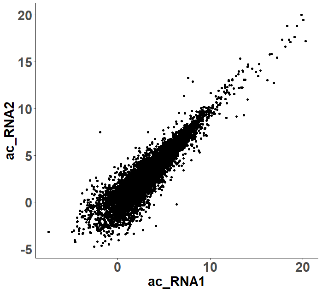

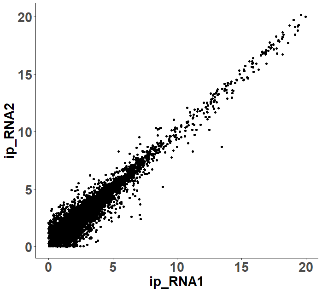

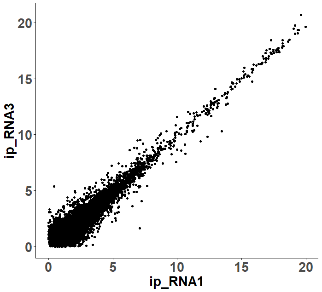

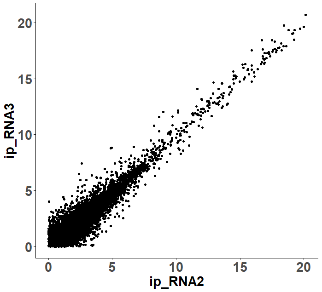

**Control_RNA1**

**Control_RNA2**

**Infection_RNA1**

**Infection_RNA2**

**Infection_RNA1**

**Infection_RNA3**

**Infection_RNA2**

**Infection_RNA3**

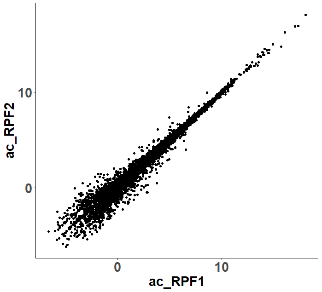

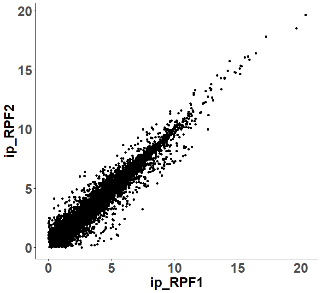

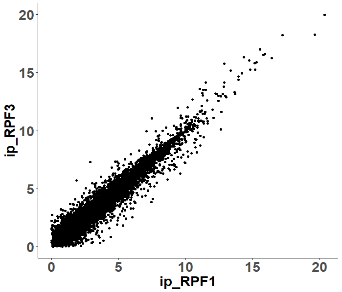

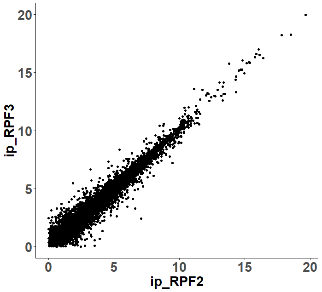

**Control_RPF1**

**Control_RPF2**

**Infection_RPF1**

**Infection_RPF2**

**Infection_RPF1**

**Infection_RPF3**

**Infection_RPF2**

**Infection_RPF3**

**r=0.91**

**r=0.98**

**r=0.96**

**r=0.95**

**r=0.95**

**r=0.95**

**r=0.96**

**r=0.97**

**Supporting information Fig. S1:** Correlation for ribo-seq and RNA-seq data within the same treatment. (a) Correlation between biological replicates of RNA samples for control and infection conditions. (b) Correlation between biological replicates of ribo-seq (RPF) samples in control and infection conditions. Correlations were determined by Pearson correlation.

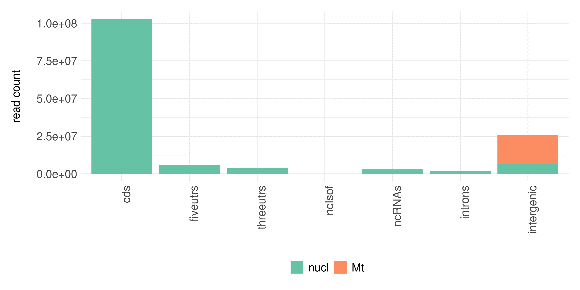

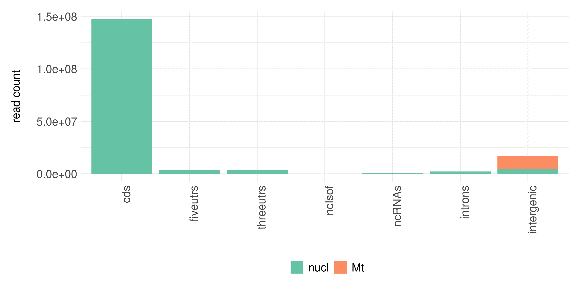

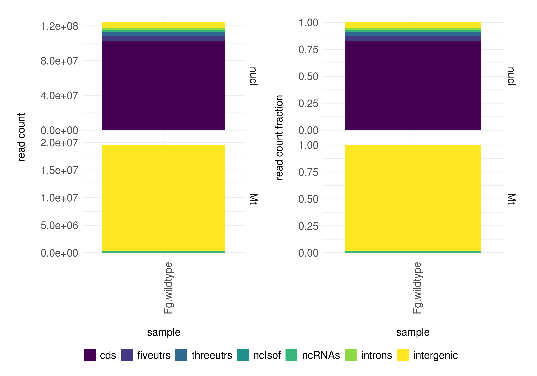

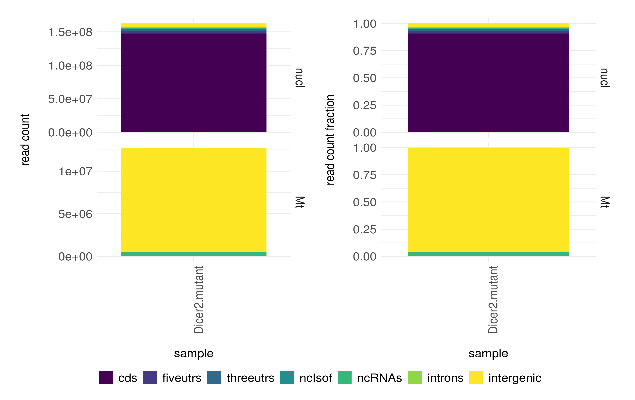

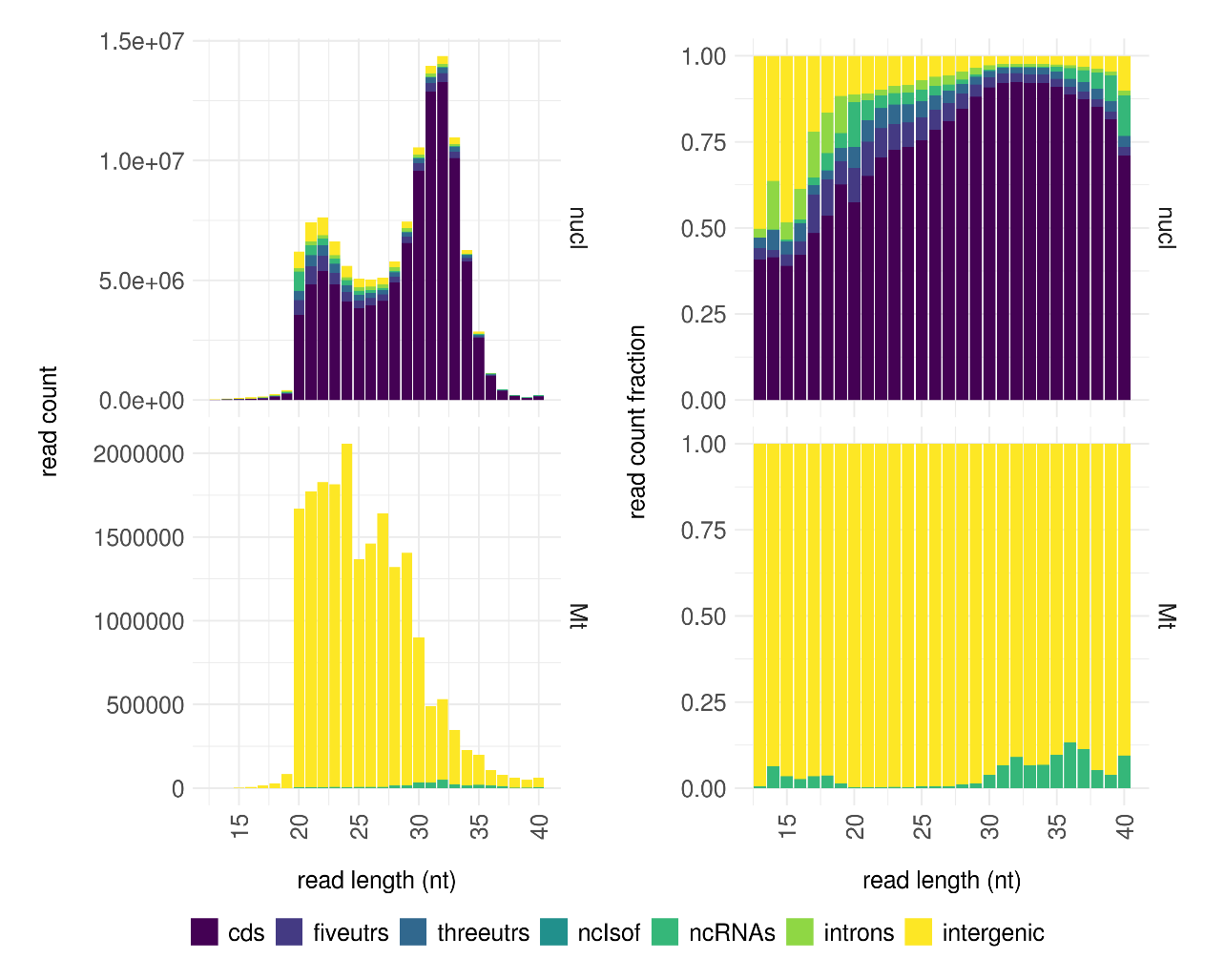

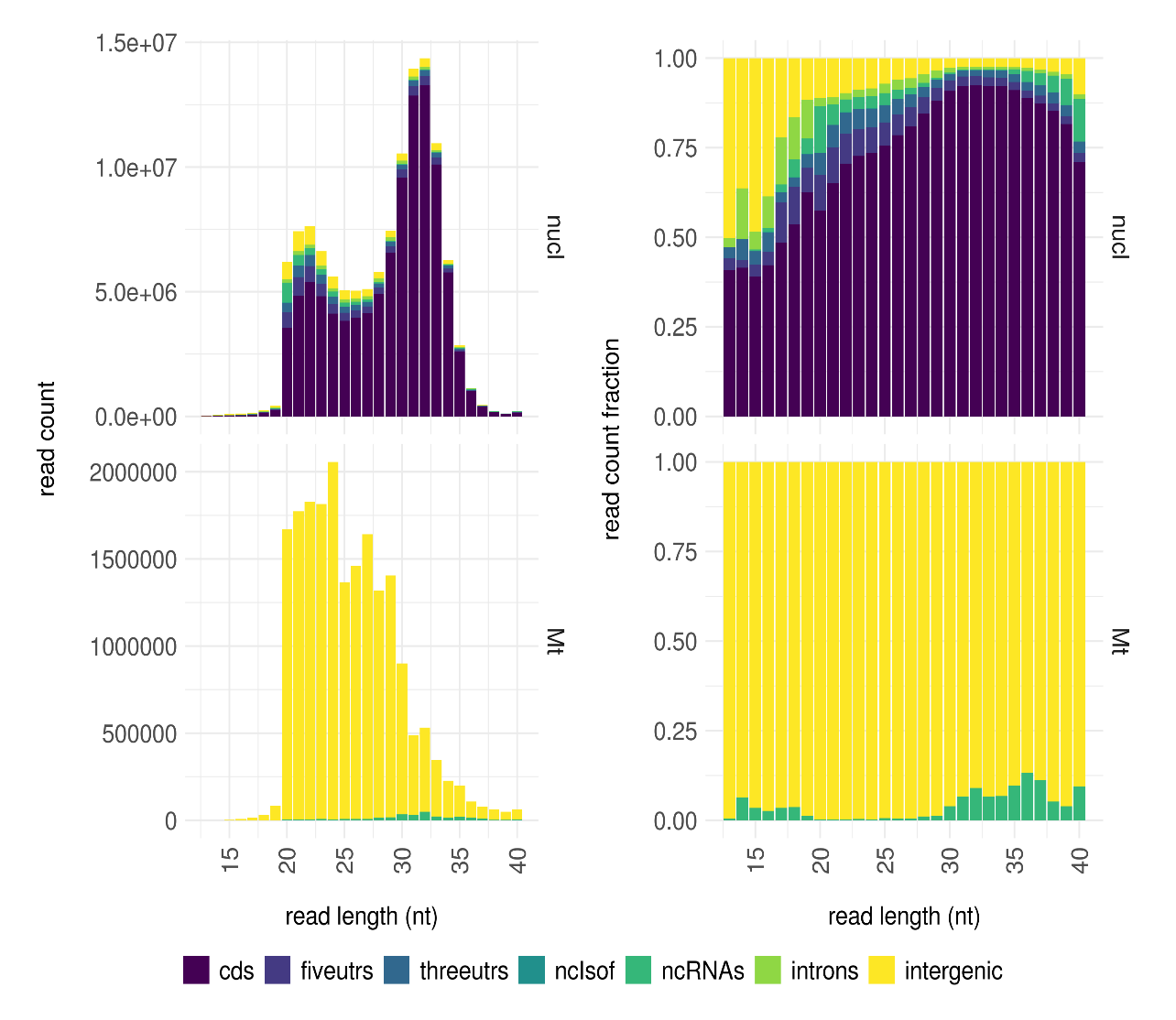

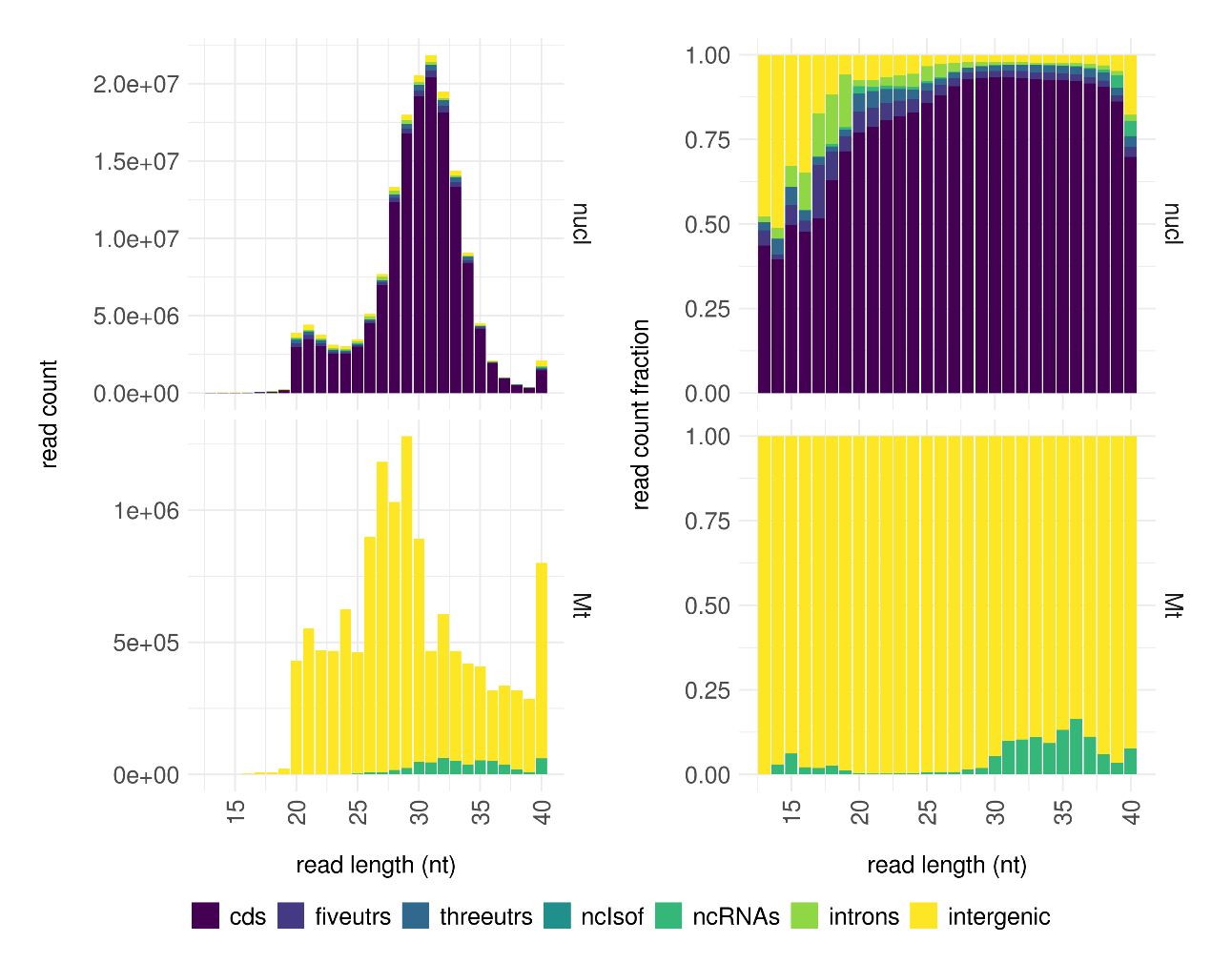

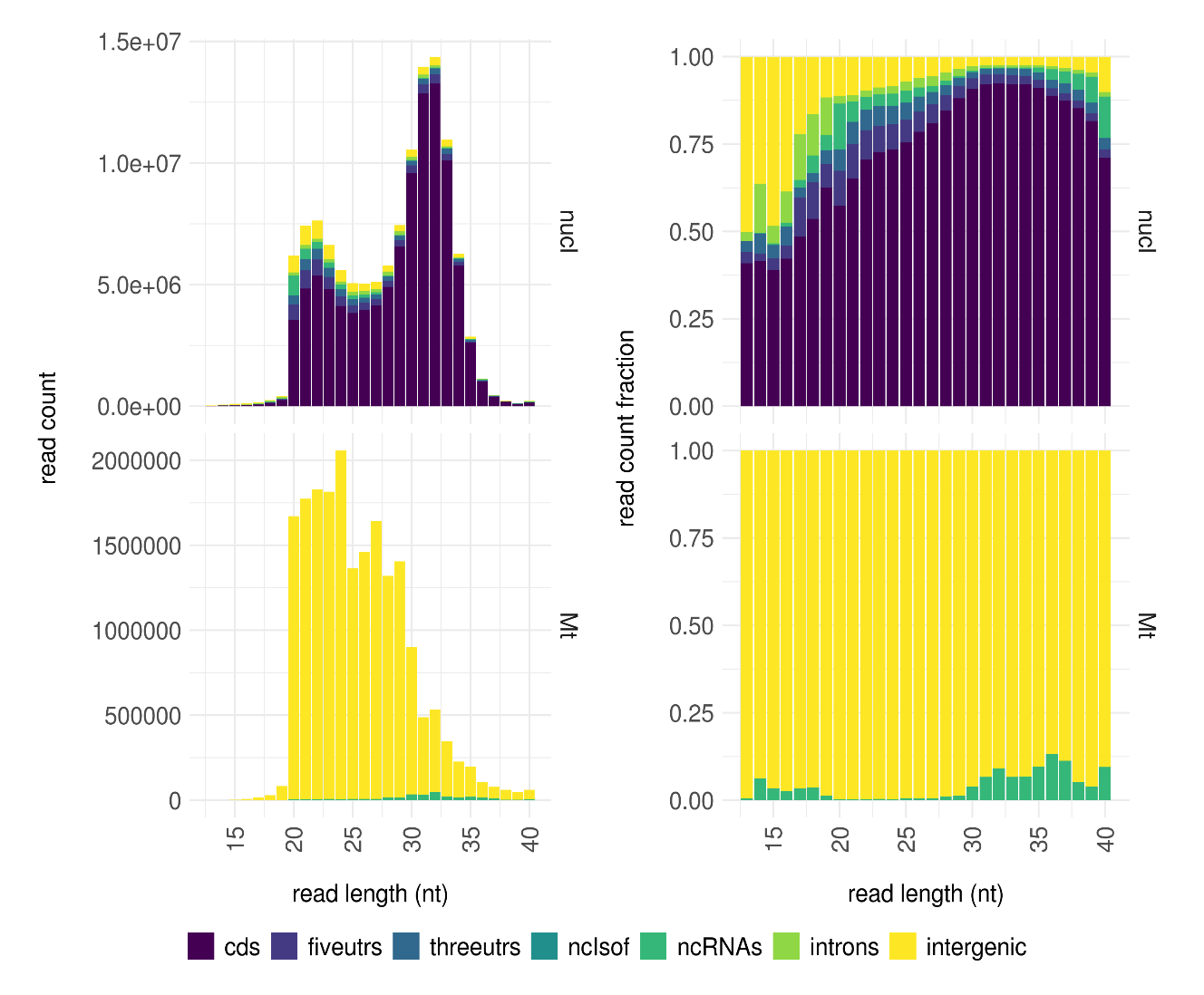

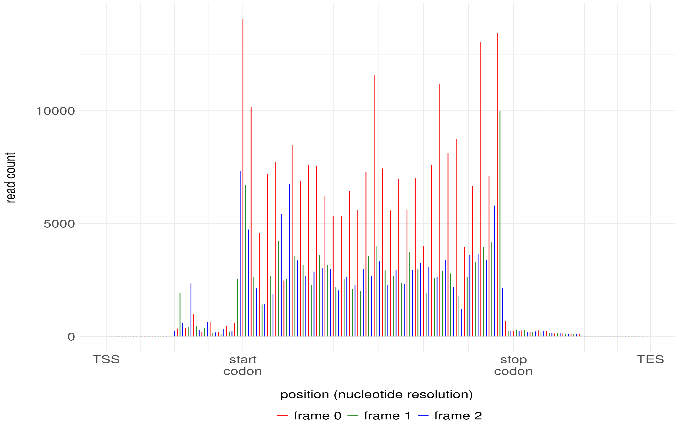

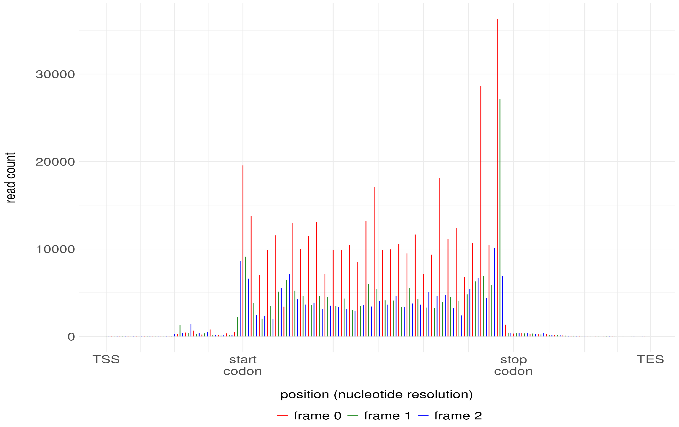

Fg_Wildtype_Ribo-seq_QC

Fg_Dicer2_Ribo-seq_QC

a)

b)

c)

**Supporting information Fig. S2:** **Ribo-seq data quality for wildtype Fg and Fgdicer2 mutant. (a)** Percent read distributions into different genomic features. **(b)** Ribosome protected footprint (RPF) read length distributions into different genomic locations. **(c)** Meta-gene analysis of 30 nt RPFs near start and stop codon showing three nucleotide periodicity.

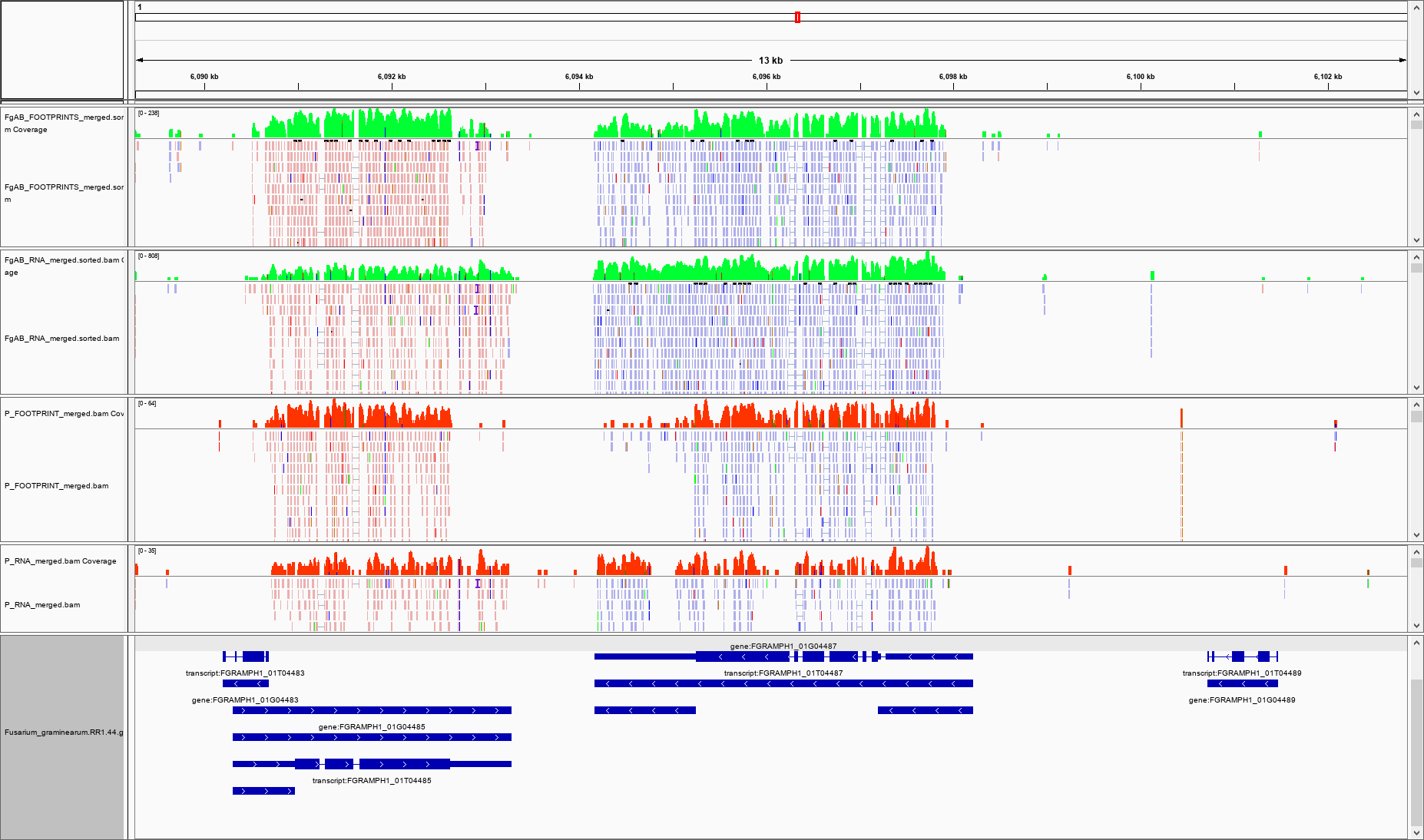

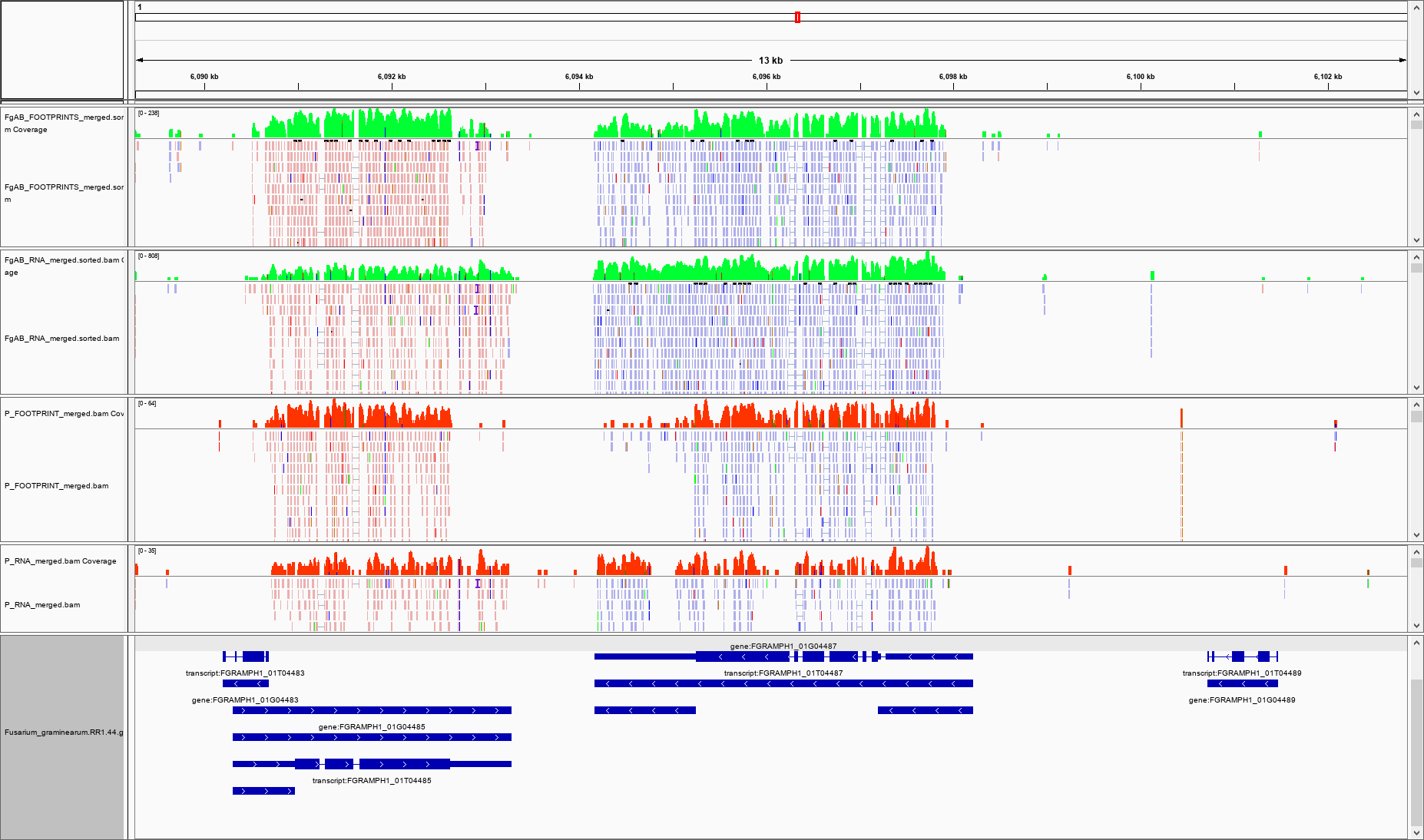

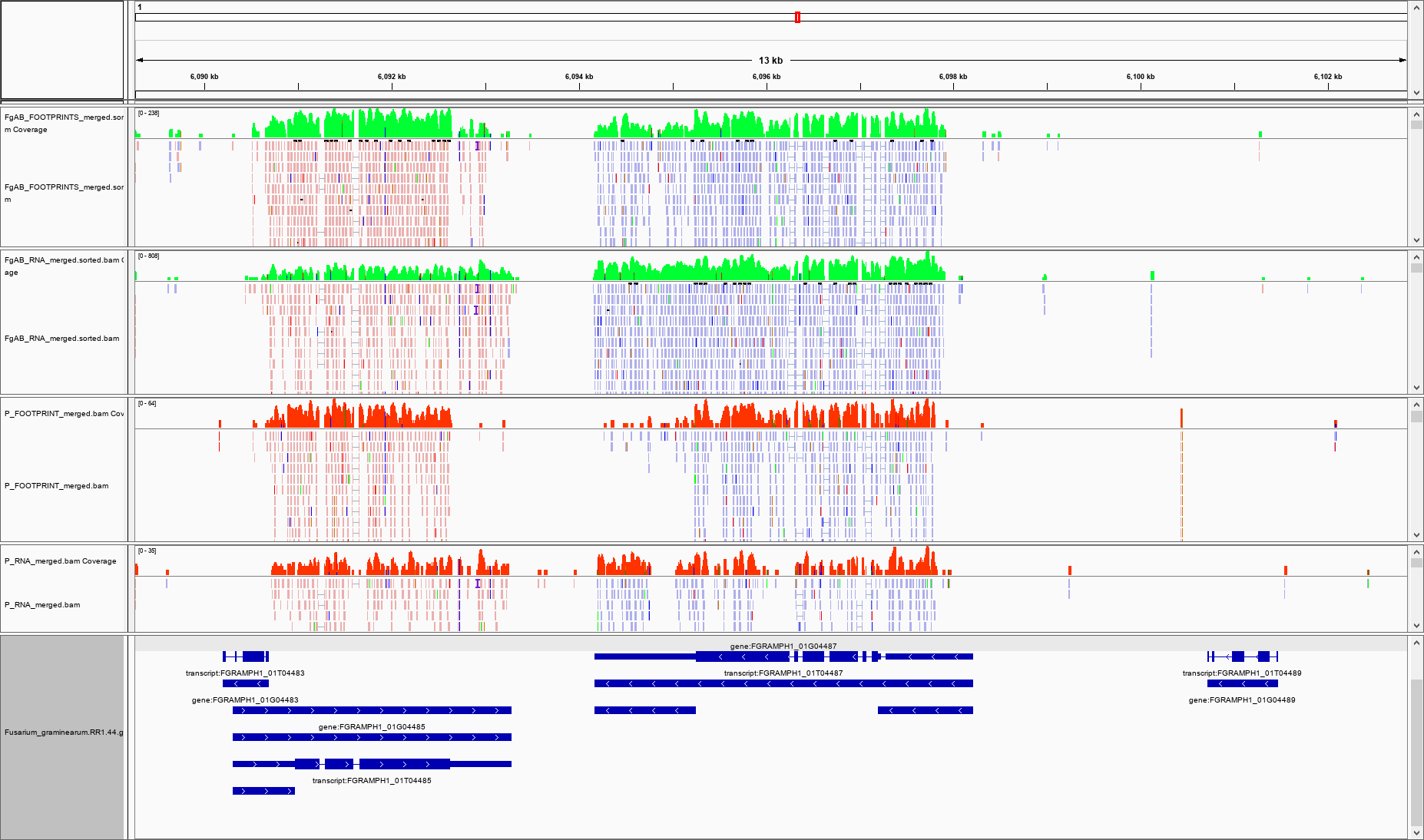

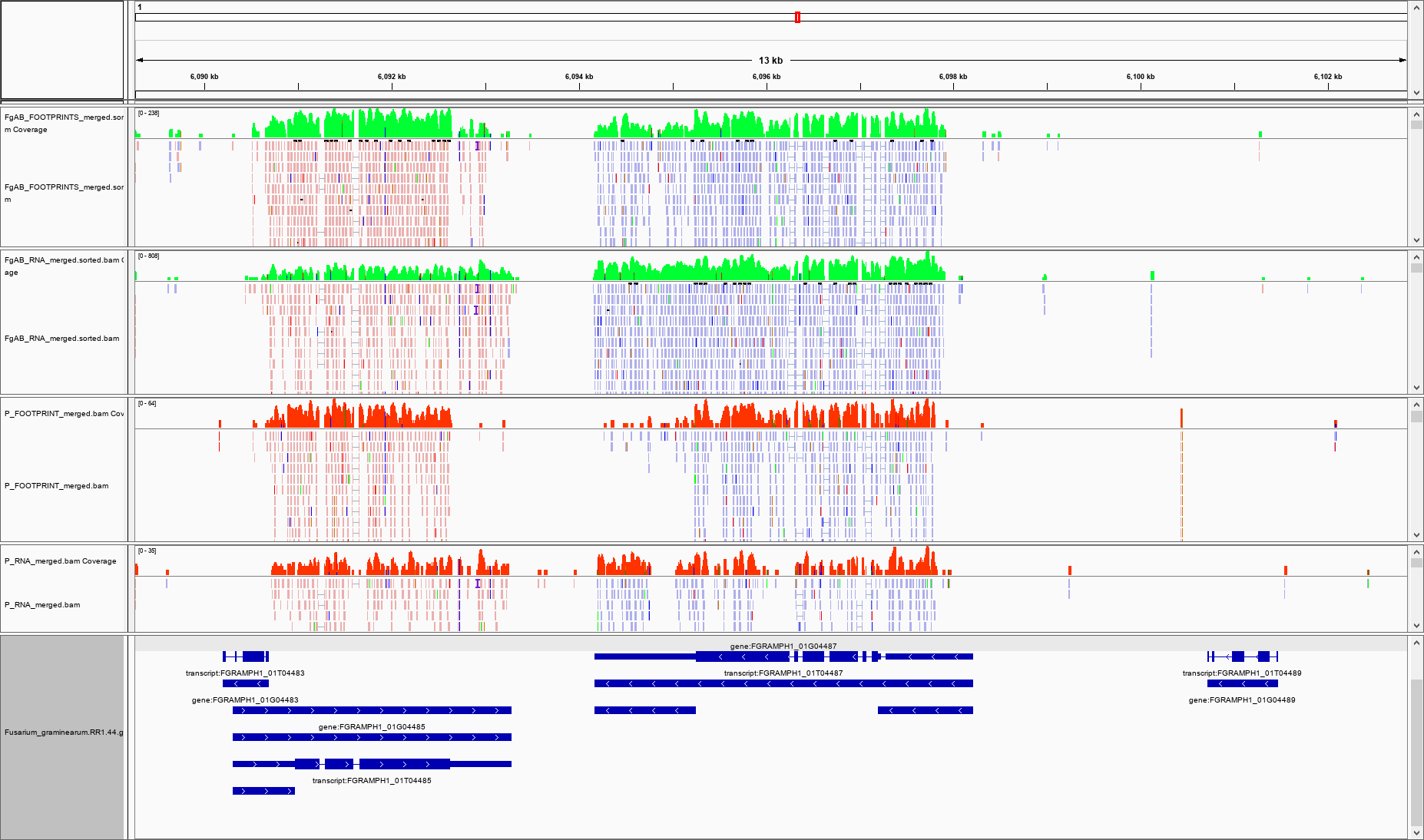

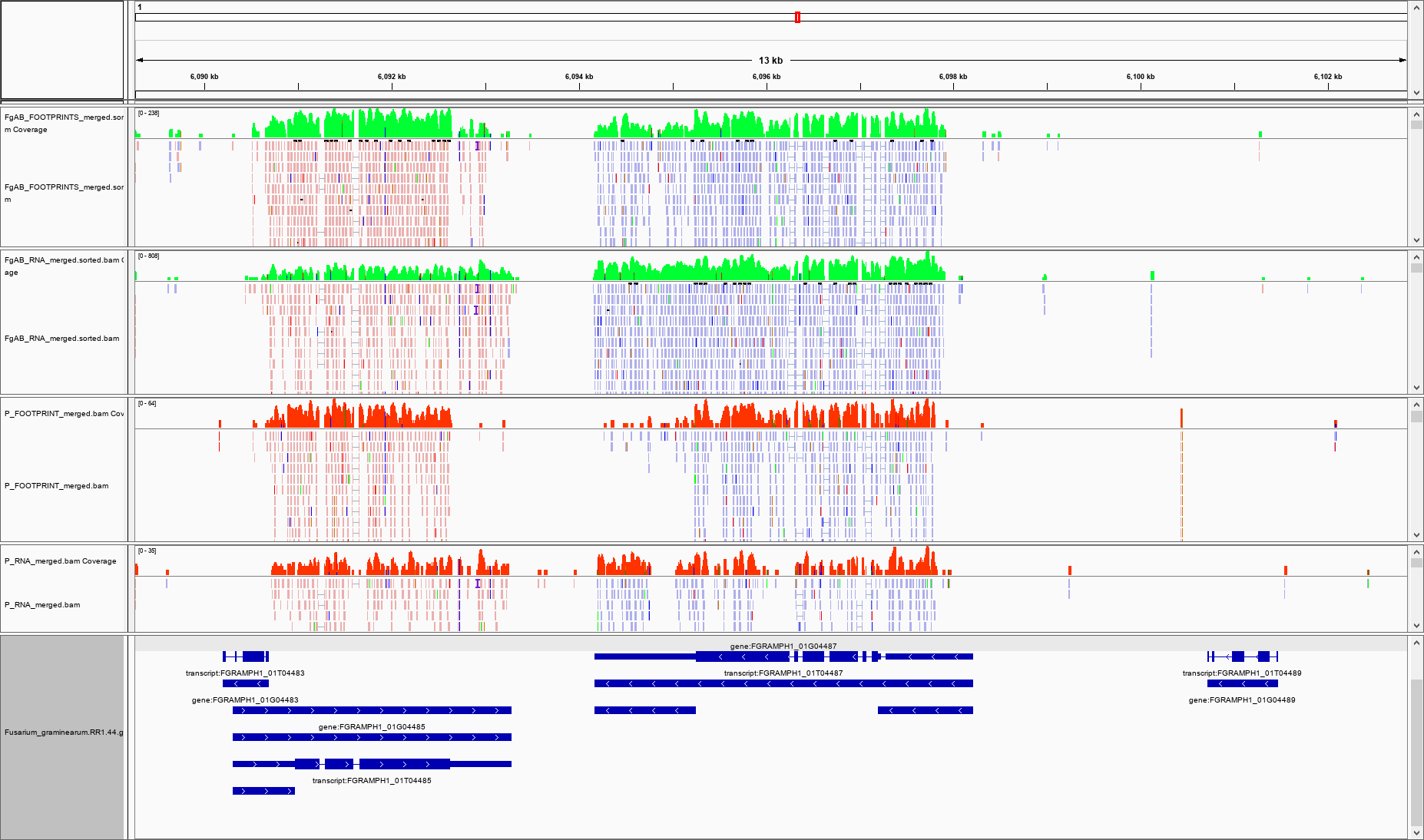

**FGRAMPH1_01T04487**

**RPF**

**RNA**

Transcript

Gene

UTRs

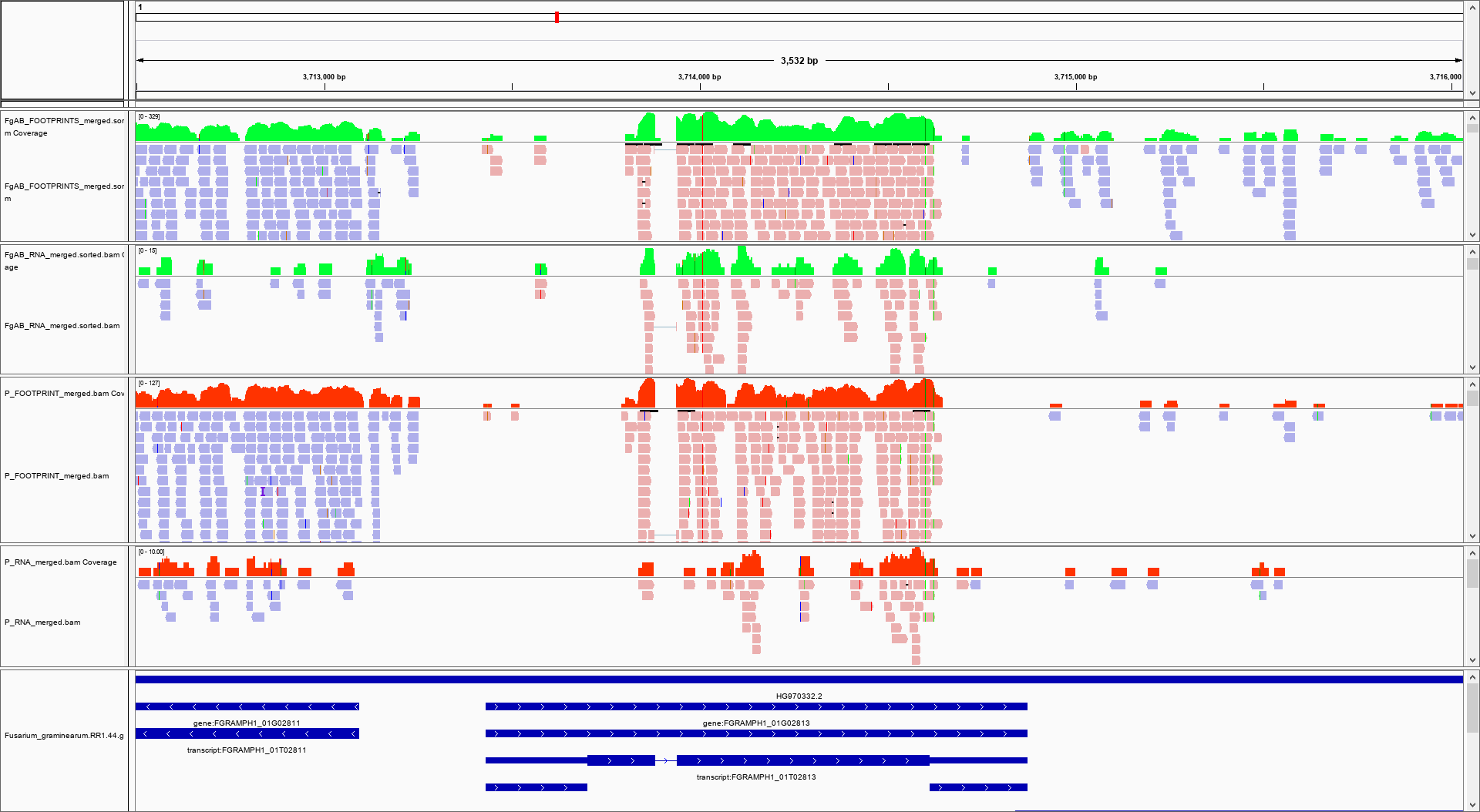

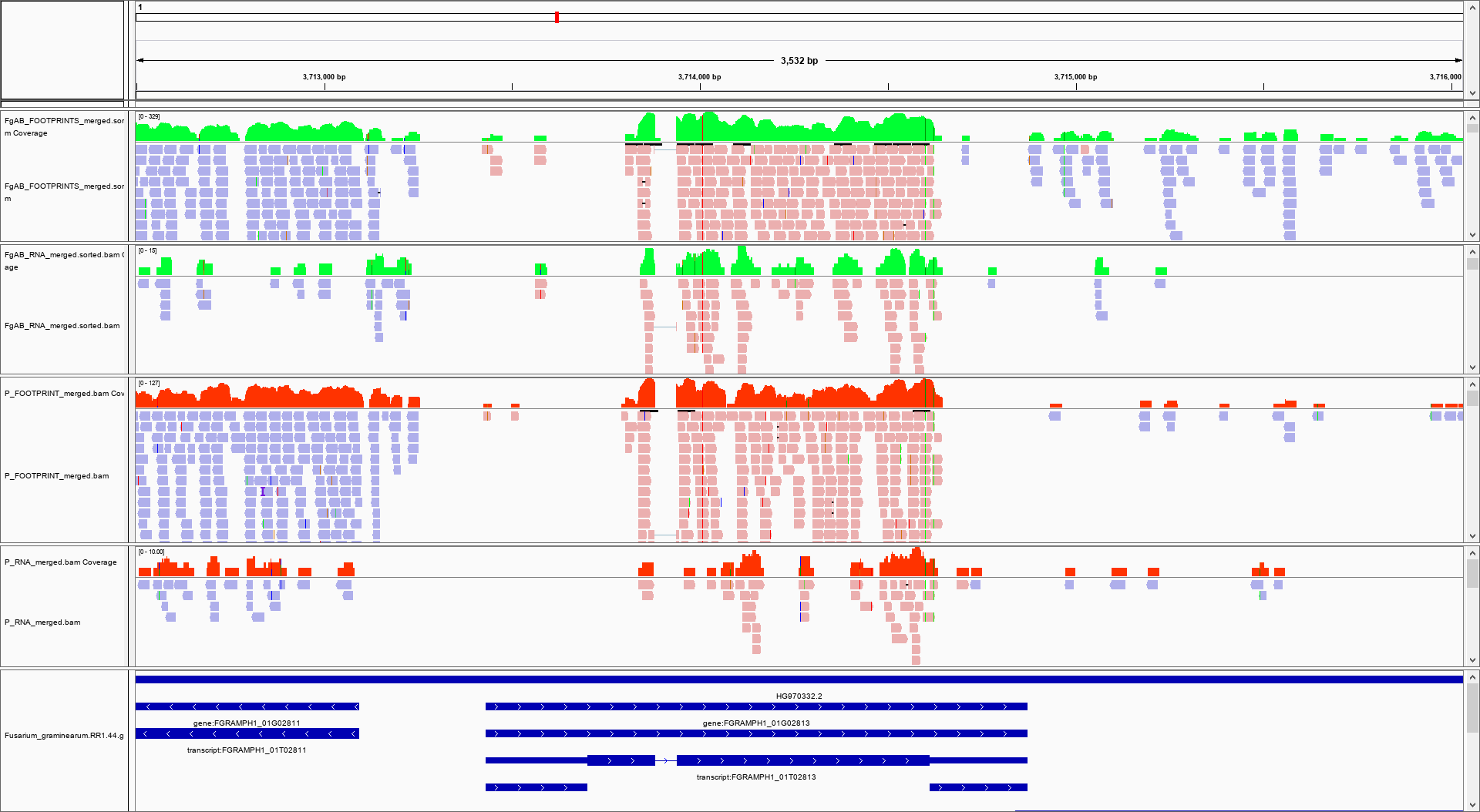

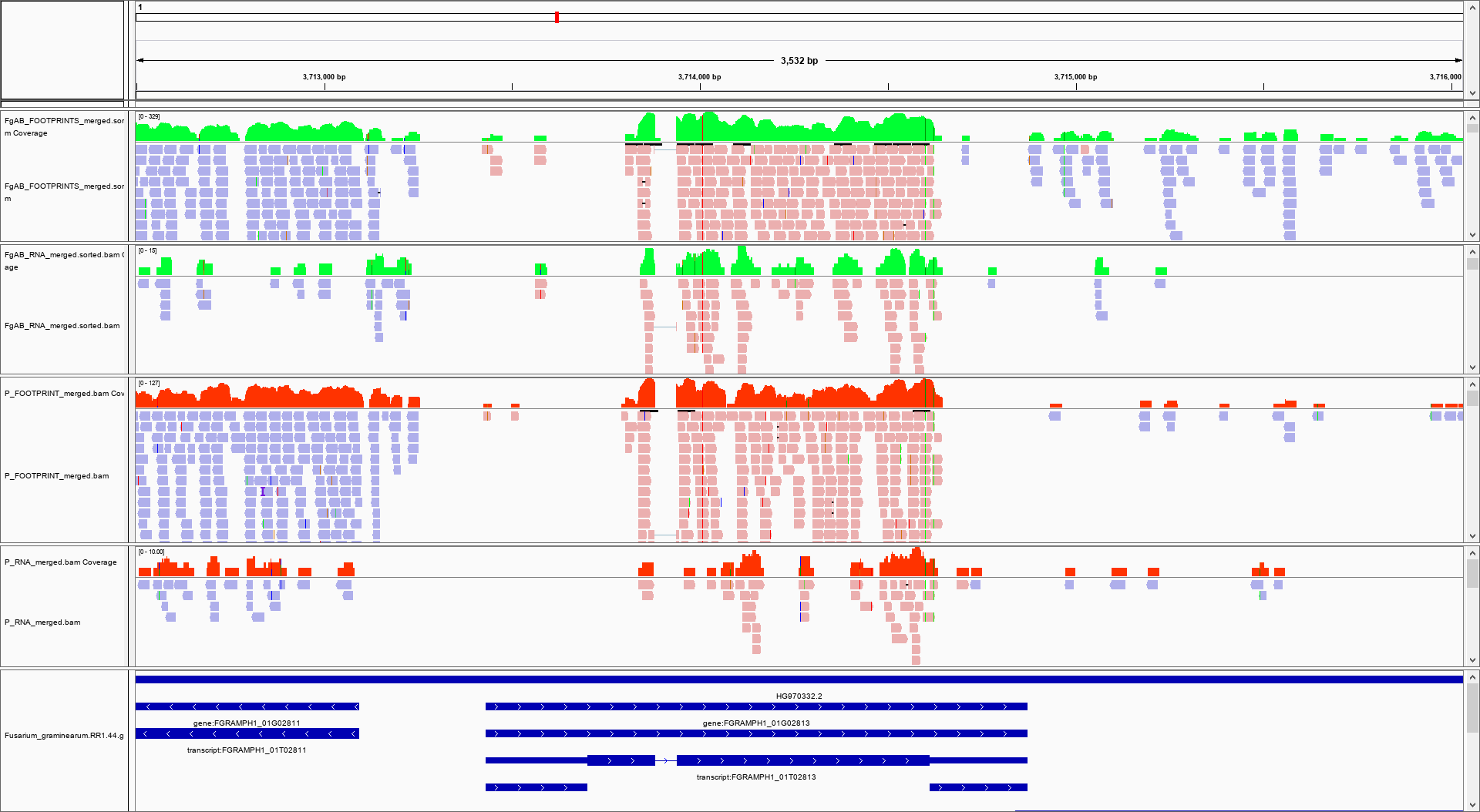

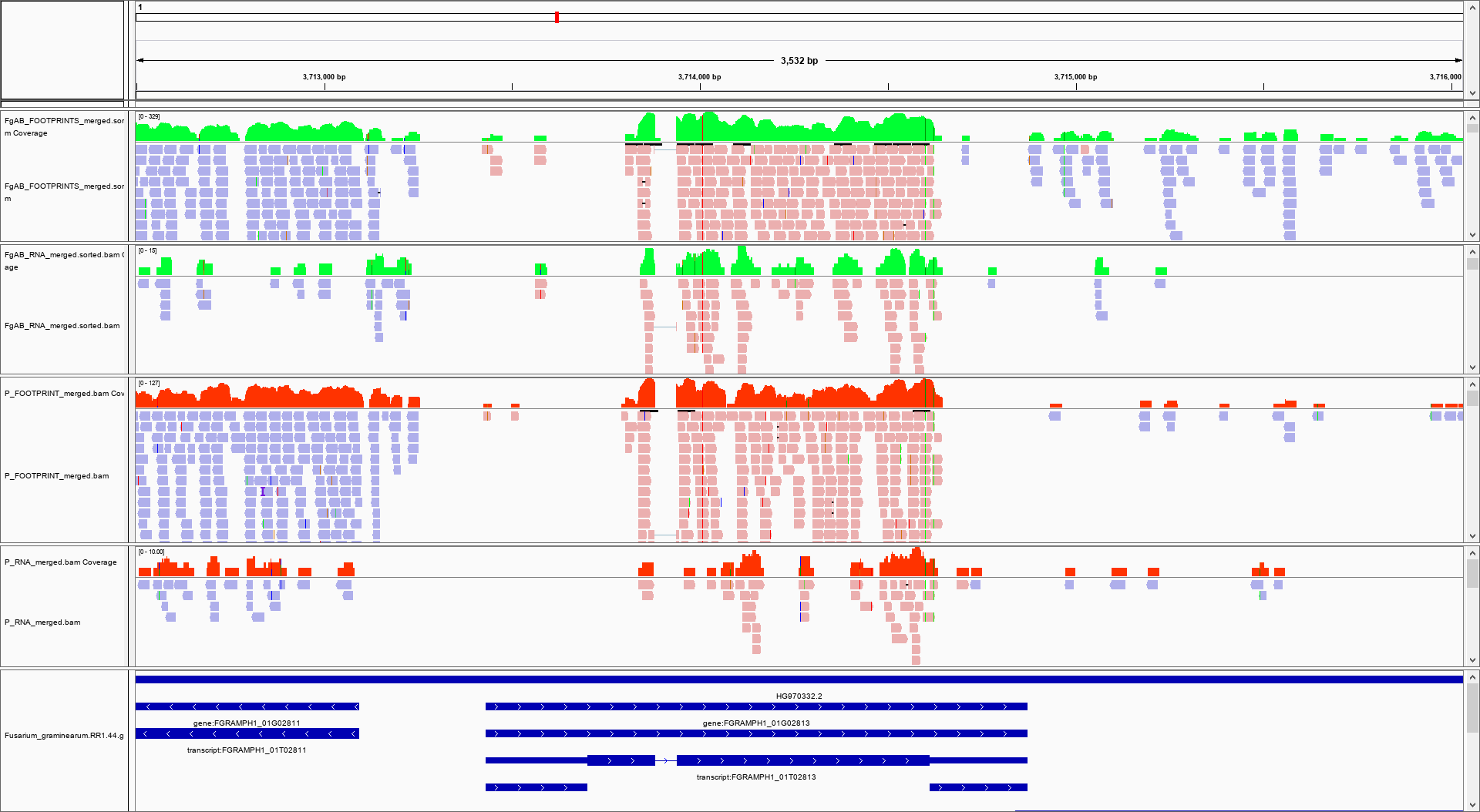

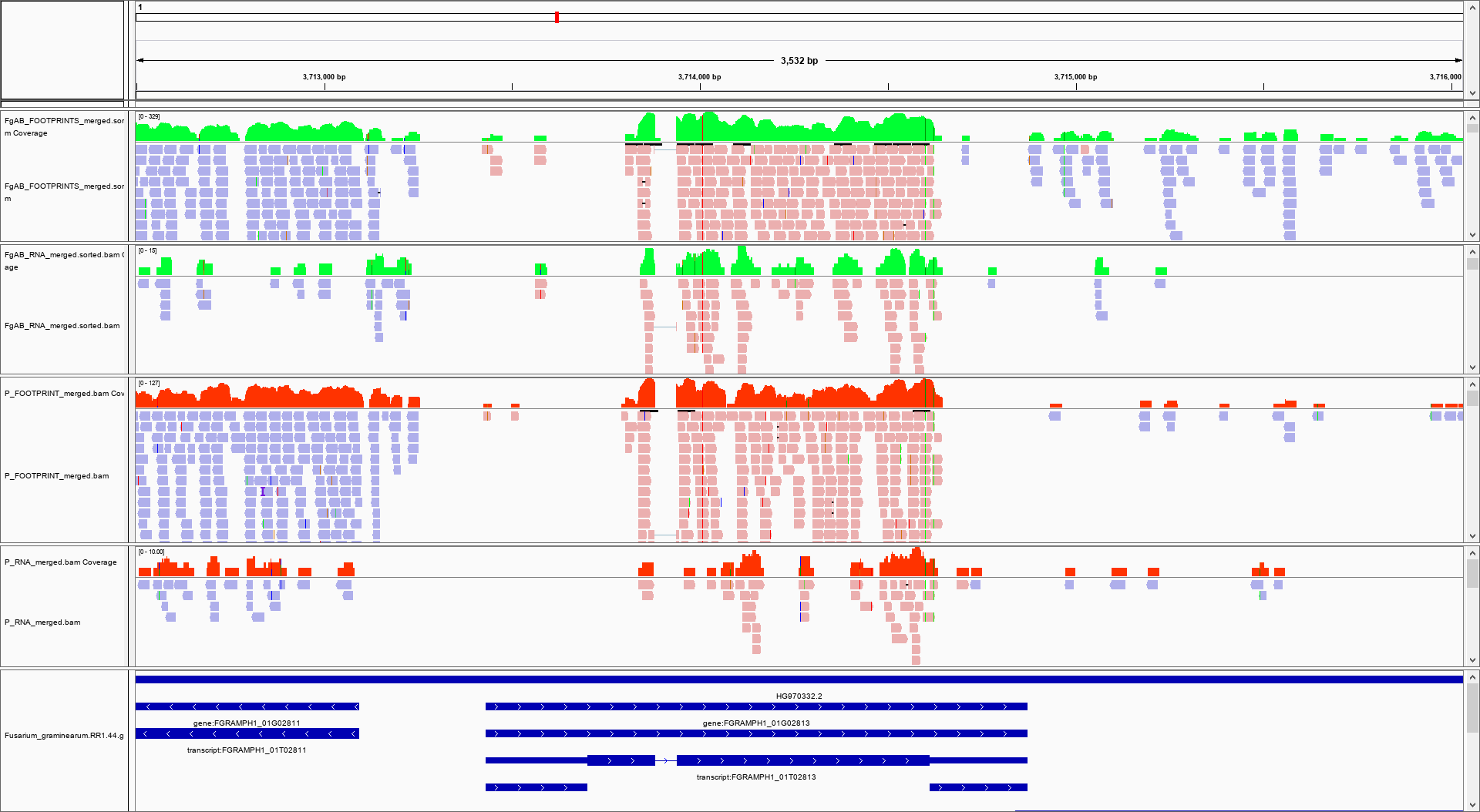

**FGRAMPH1_01T02813**

**RPF**

**RNA**

**a**

**b**

**Supporting information Fig. S3: Novel ORFs identified from ribo-seq data in *Fg.* (a)** Ribo-seq data shows the ORF translating from upstream to the annotated start site and running down to the annotated stop site. **(b)** Possible downstream start of the ORF from the annotated start site. Black and red dashed lines indicate annotated start and stop sites respectively.

FgNir 5’ Flank

**Nat1**

FgNir 5’ Flank

**FgNir**

FgNir 3’ Flank

FgNir 3’ Flank

**Nat1**

Nat1 terminator

Primer1F

Primer2R

Nat1 terminator

Primer1R

Primer2F

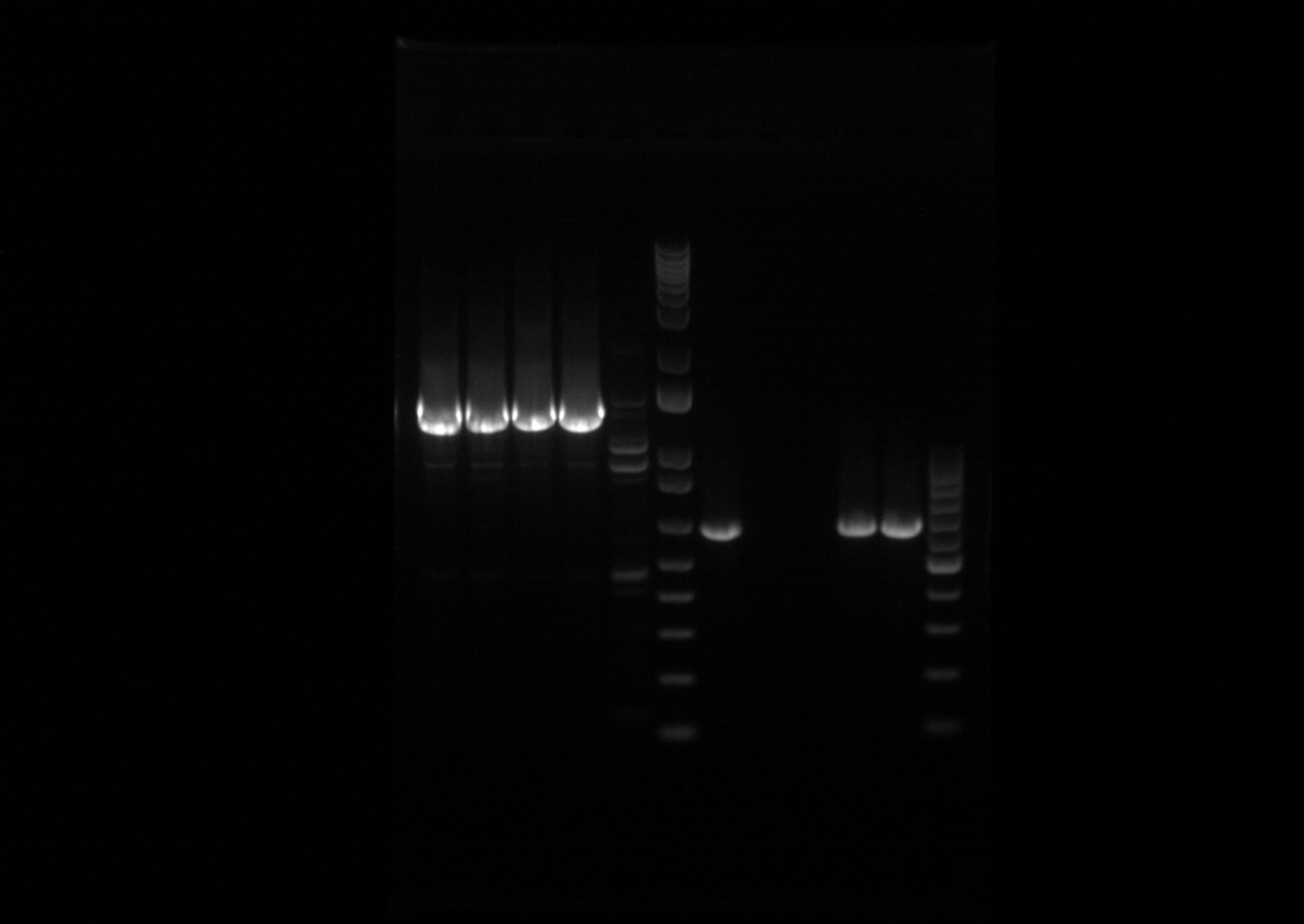

1

2

3

4

Wt-Fg

100bp+

1kb+

~600bp

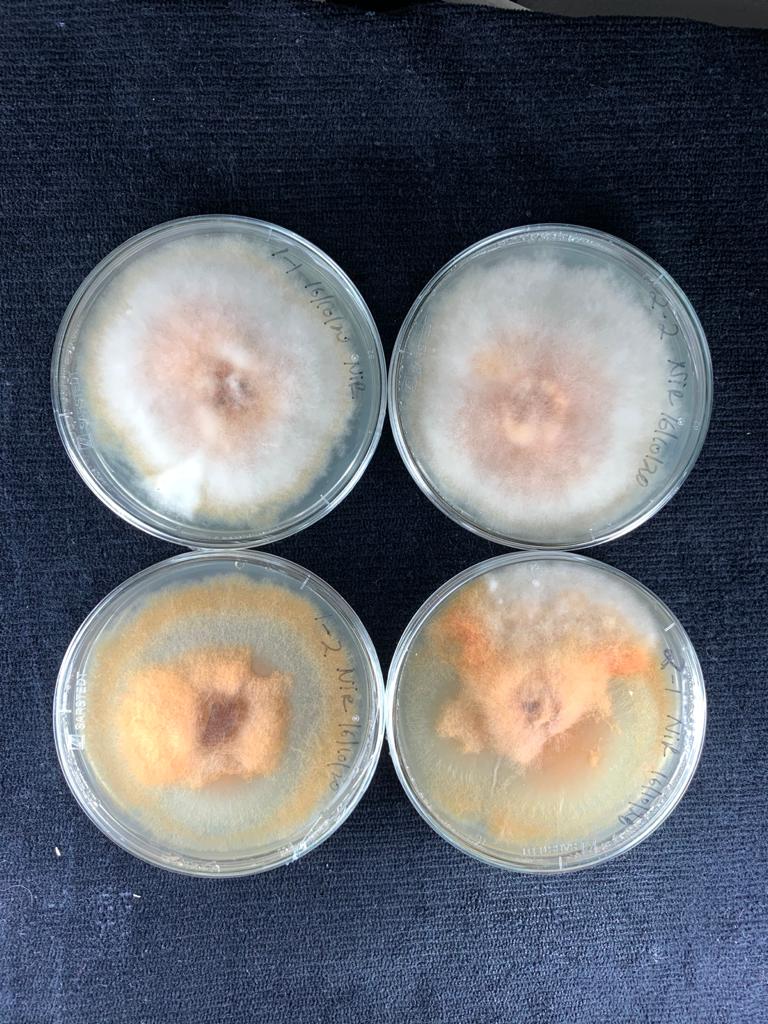

**Nitrite reductase amplification**

**using primer pair 1**

1

4

2

3

**Supporting information Fig. S4:** FgNir mutant development using split marker strategy and mutant screening by PCR. Mutant were confirmed with primers pairs 1 and 2 (data not shown). Number 1-4 in gel pictures are different transformants and Wt-Fg is wild type *F. graminearum*. Right hand side picture represents phenotypes of transformants on the PDA plates.

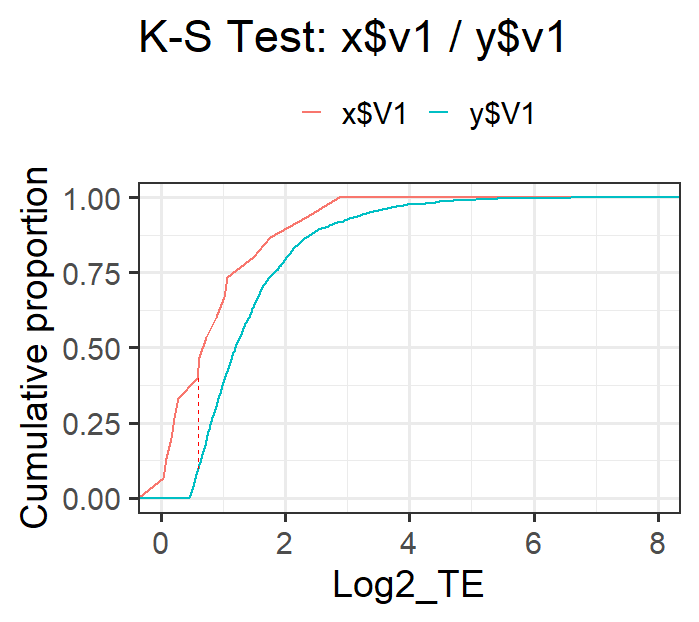

Genes with translating uORF

Genes without translating uORF

p=0.04314

**ECDF**

**Log2_TE**

D=0.35906

**Supporting information Fig. S5: Effect of upstream ORFs** **(uORF) on translational efficiency of mORF under control condition in *Fg.*** Red dotted line is absolute difference (D) between the two distributions called as K-S statistic. The p and D-values were determined with two-sample Kolmogorov-Smirnov test.
